# PDAC BioPortal: An Integrated Web Resource for Multi-Scale Analysis of Pancreatic Cancer Transcriptomics

**DOI:** 10.64898/2026.11.18.767683

**Authors:** Prince Kumar, Erina Vlashi

## Abstract

Pancreatic ductal adenocarcinoma (PDAC) exhibits profound cellular heterogeneity and treatment-associated remodeling, yet bulk, single-nucleus, and spatial transcriptomic datasets remain fragmented across separate portals. We developed PDAC BioPortal, an open-access, browser-based web resource integrating four complementary human PDAC cohorts: primary tumor versus normal pancreas (TCGA-PAAD and GTEx, n = 345), a neoadjuvant stereotactic body radiotherapy-containing bulk cohort (GSE225767, n = 55), a single-nucleus chemoradiotherapy atlas (GSE202051, 224,988 nuclei across 43 patients), and 10x Genomics Visium spatial transcriptomics (GSE274103, 23,436 spots across five tumors). The platform provides interactive differential expression, rank-based pathway enrichment, patient-level pseudobulk comparisons, survival exploration, and spatial visualization, alongside PDACopilot, a deterministic, evidence-grounded natural-language query interface. Across cohorts, neoadjuvant-treated bulk and single-nucleus pathway rankings showed significant concordance (Spearman ρ = 0.512, p = 1.46 × 10⁻⁴), while Kitagawa-Oaxaca-Blinder decomposition and compositional sensitivity testing separated epithelial debulking from cell-intrinsic signaling shifts. Interrogation of the NRF2-associated redox network illustrated how bulk changes diverge from cell-type-resolved and spatial patterns, while continuous Cox modeling linked higher NRF2-signature expression to shorter overall survival. PDAC BioPortal is freely available at https://limboper89.github.io/PAAD-SBRT-GEx-Dashboard/.

Availability: https://limboper89.github.io/PAAD-SBRT-GEx-Dashboard/

## 1. Introduction

Pancreatic ductal adenocarcinoma (PDAC) remains one of the most lethal and treatment-refractory human malignancies, characterized by profound biological heterogeneity and an exceptionally dense, hypovascular fibroinflammatory stroma ^[1,2]^. Disease progression is governed not only by cell-intrinsic oncogenic drivers in malignant epithelial cells, predominantly activating KRAS mutations and loss of TP53, CDKN2A, and SMAD4 ^[3]^, but also by dynamic reciprocal cross-talk with cancer-associated fibroblasts (CAFs), vascular endothelium, infiltrating myeloid lineages, and immunosuppressive lymphoid populations ^[4,5]^. This extensive desmoplasia imposes severe physical and metabolic constraints, driving regional tissue hypoxia, nutrient deprivation, and resistance to standard systemic therapies ^[1,6]^. Crucially, the PDAC tumor ecosystem is not biologically static: neoadjuvant interventions, including multi-agent chemotherapy and radiotherapy substantially remodel both the epithelial and stromal compartments, generating complex transcriptional, metabolic, and architectural states that diverge fundamentally from those observed in untreated tumors ^[5,7,8]^. Consequently, deciphering the mechanisms that govern therapeutic resistance requires systematic interrogation across multiple biological scales, spanning whole-tissue alterations, single-cell heterogeneity, spatial neighborhoods, and treatment-associated responses across disease stages.

Recent advances in high-throughput sequencing have generated a rich landscape of publicly available PDAC transcriptomic datasets that capture these complementary dimensions. Population-scale resources, such as The Cancer Genome Atlas (TCGA-PAAD) and the Genotype-Tissue Expression (GTEx) project, provide foundational references for cataloging molecular alterations that distinguish untreated primary tumors from non-malignant pancreas ^[9]^. Subsequently, single-cell and single-nucleus RNA sequencing (snRNA-seq) studies have deconvolved the cellular diversity of the tumor microenvironment, revealing transcriptionally distinct CAF subtypes (including inflammatory iCAFs and myofibroblastic myCAFs), heterogeneous endothelial states, and diverse immune phenotypes ^[4,5,10]^. More recently, spatially resolved transcriptomics has enabled functional gene programs to be localized within their native histological niches ^[6]^, while clinically annotated treatment cohorts have begun to capture the molecular remodeling that accompanies neoadjuvant therapy ^[5,7]^.

The growing volume of these datasets has spurred the development of valuable platforms designed to facilitate their exploration. Portals such as GEPIA2 ^[11]^ enable rapid interactive queries across TCGA and GTEx cohorts, while cBioPortal ^[12,13]^ provides broad access to multidimensional genomic, copy-number, and clinical cancer profiles. In the single-cell arena, resources such as TISCH2 ^[14]^ and CELLxGENE ^[15]^ have standardized access to tumor microenvironmental scRNA-seq datasets across multiple cancer types, while specialized platforms such as TIGER ^[16]^ integrate bulk and single-cell data to interrogate tumor immunogenicity and immunotherapy responses.

Despite these technological and computational strides, a critical analytical gap persists - public PDAC datasets remain fragmented across isolated portals and experimental modalities. Bulk, single-nucleus, spatial, and treatment-associated datasets are typically generated by independent consortia and queried in isolation. This separation is particularly consequential in PDAC because the biological interpretation of a molecular signature depends fundamentally on the scale at which it is evaluated ^[5,17]^. In whole-tissue bulk RNA-seq, an apparent change in gene or pathway expression frequently reflects alterations in tissue composition, such as therapy-induced parenchymal debulking and fibroblastic expansion-rather than cell-autonomous transcriptional reprogramming ^[5,17]^. Conversely, critical cell-type-restricted survival adaptations may be masked or heavily diluted when averaged across a dense desmoplastic background ^[17]^. Without an integrated environment that enables continuous movement between bulk cohorts, cellular compartments, and spatial architectures, researchers risk misattributing tissue-level compositional shifts to cell-intrinsic mechanisms or overlooking localized niches of therapeutic resistance.

This fragmentation is especially acute in the context of radiation therapy. While stereotactic body radiotherapy (SBRT) and hypofractionated radiation are increasingly incorporated into neoadjuvant regimens for borderline-resectable and locally advanced PDAC ^[7,18,19]^, transcriptomic data from irradiated human PDAC tissues remain scarce and technically difficult to isolate within primary repositories such as the Gene Expression Omnibus (GEO). Furthermore, existing radiation-associated datasets (such as unpaired pre-treatment biopsies and post-neoadjuvant surgical resections; ^[7]^) have remained siloed in study-specific repositories, unintegrated with single-cell or spatial references that can resolve which cell types drive post-radiation remodeling. Given that ablative radiation triggers profound simultaneous changes across malignant cells, the microvasculature, immune infiltrates, and the extracellular matrix ^[7,10]^, the lack of an integrated comparative platform has impeded a comprehensive understanding of radiation-associated tissue remodeling in PDAC.

To bridge these gaps, we developed PDAC BioPortal, an open-access analytical framework integrating four complementary PDAC transcriptomic resources: primary tumors versus normal pancreas (TCGA-PAAD and GTEx) ^[9,20]^, a neoadjuvant bulk cohort incorporating chemotherapy and SBRT ^[7]^, a treatment-stratified single-nucleus atlas ^[5]^, and 10x Genomics Visium spatial transcriptomics ^[21]^. The platform supports interactive differential expression, over-representation analysis (ORA), pre-ranked gene set enrichment analysis (GSEA), patient-aware pseudobulk comparisons, survival exploration, and spatial visualization, alongside PDACopilot, an evidence-grounded conversational assistant ^[22,23]^. Using this platform, we examined cross-cohort pathway concordance, resolved compositional versus cell-intrinsic shifts, and interrogated the NRF2-associated redox network as a biological case study across disease and treatment contexts.

## 2. Results

### 2.1. Overview of PDAC BioPortal

PDAC BioPortal harmonizes four complementary transcriptomic cohorts across disease, treatment, cellular, and spatial scales (Figure 1; Table 1). At the gene and cohort level, investigators can interrogate baseline tumor-versus-normal expression, treatment-associated bulk differences, pathological response classifications, and clinical survival associations with interactive statistical summaries and downloadable outputs (Table 2).

**Figure 1.**
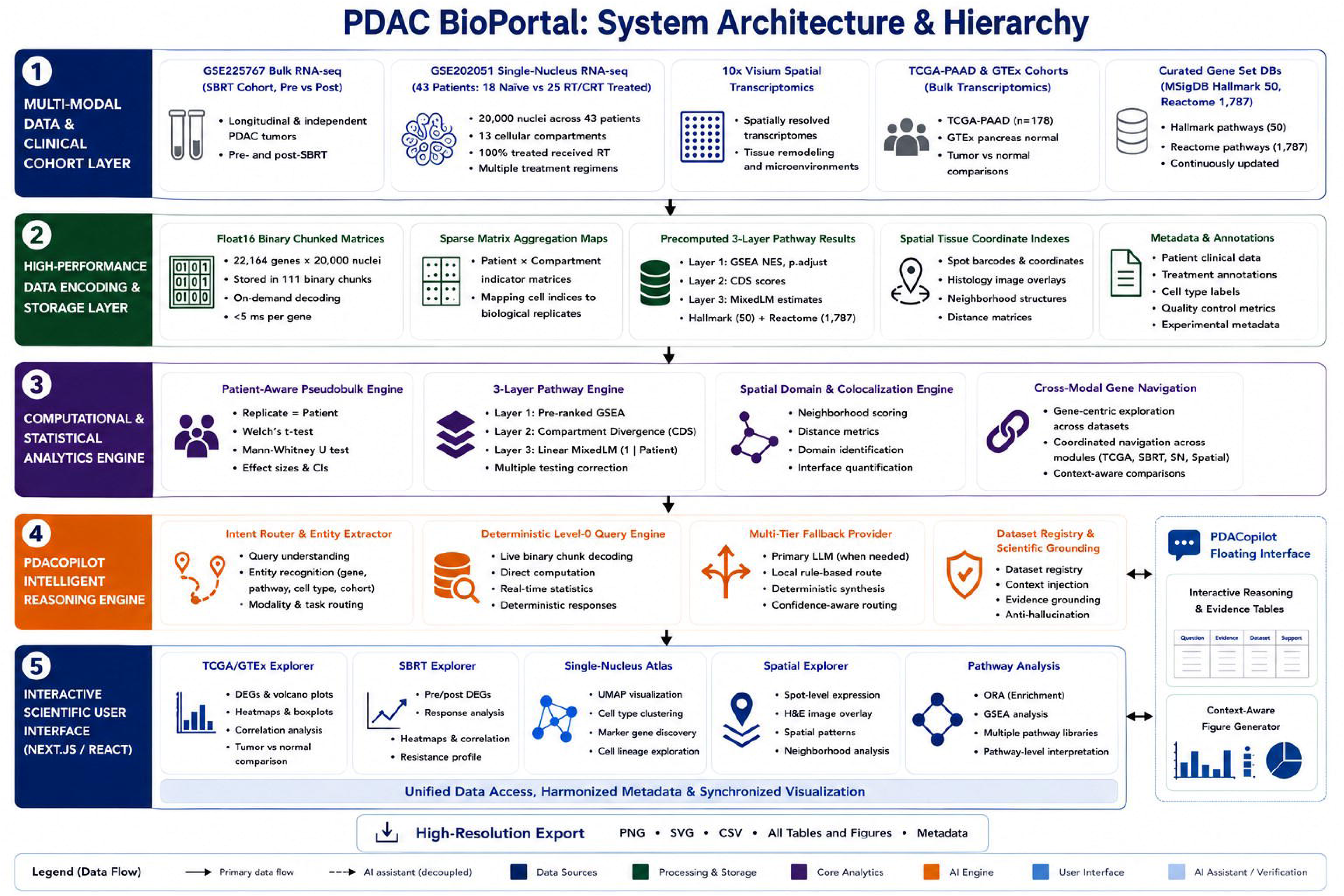
System architecture and multi-modal analytical workflow of PDAC BioPortal. PDAC BioPortal integrates four complementary human pancreatic ductal adenocarcinoma (PDAC) transcriptomic cohorts (TCGA-PAAD primary tumors [n = 178] vs. GTEx normal pancreas [n = 167] and TCGA solid-tissue normal controls [n = 4], GSE225767 neoadjuvant SBRT bulk RNA-seq [n = 26 pre-SBRT vs. n = 29 post-SBRT], GSE202051 single-nucleus RNA-seq [224,988 nuclei across 43 patients; 20,000-nucleus stratified interactive subset], and GSE274103 10x Genomics Visium spatial transcriptomics [23,436 spots across 5 patients]) with curated MSigDB Hallmark (50), Reactome (1,787), and GO Biological Process (5,393) pathway collections. Web-optimized Float16/Float32 binary matrices, patient-level pseudobulk aggregation, spatial coordinate stores, and the three-tier PDACopilot reasoning engine power synchronized interactive modules with deterministic PNG, SVG, PDF, and CSV export.

**Table 1.** Summary of transcriptomic datasets integrated into PDAC BioPortal.

| **Dataset** | **Samples** | **Technology** | **Primary application** |
|---|---|---|---|
| Tumor vs. normal | 349 portal samples (primary DE: 178 TCGA tumors vs. 167 GTEx normal pancreas, n = 345; plus 4 TCGA solid-tissue normal controls) | Bulk RNA-seq | Tumor-versus-normal differential expression and correlation analyses |
| Pre- vs. post-SBRT | 55 PDAC samples: 26 pre-SBRT, 29 post-SBRT | Bulk RNA-seq | SBRT-associated transcriptomic changes |
| Single-nucleus RNAseq | 224,988 nuclei 43 patients (18 treatment-naïve; 25 neoadjuvant-treated). | Single-nucleus RNA-seq | Cell-type-specific gene expression in treatment naive vs Neoadjuvant treated/CRT |
| Spatial transcriptomics | 23,436 spatial spots; 5 patients-treatment naive tumors | 10x Genomics Visium | Spatial gene expression and tissue architecture |

**Table 2.** Key features of the PDAC BioPortal platform.

| **Feature** | **Supported** |
|---|---|
| Interactive visualization | Yes |
| Differential expression | Yes |
| Correlation analysis | Yes |
| Heatmap generation | Yes |
| Publication-quality figure export | Yes |
| CSV export | Yes |
| Cell metadata export | Yes |

By linking bulk cohorts with the treatment-stratified single-nucleus atlas, users can filter patients by neoadjuvant regimen, compare treatment subgroups against treatment-naïve controls at lineage or sub-lineage resolution using patient-aware pseudobulk aggregation, and evaluate pathway enrichment across whole-atlas and compartment-resolved profiles. Beyond precomputed cohorts, the pathway module supports interactive ORA and rank-sum GSEA (plus custom user gene-list uploads) with enrichment curves, leading-edge overlap matrices, and pathway network graphs.

Static deployment optimization reduced the asset footprint from 112,012 to 567 files, yielding initial page-load times under 350 ms and indexed gene queries in 10-30 ms in browser memory without server-side computation. Together, these modules establish a continuous workflow from gene-level differential expression to pathway networks, cellular compartments, and spatial tissue architecture.

Collectively, PDAC BioPortal establishes a unified framework in which molecular observations can be followed from individual genes and selected gene sets to pathway-level analyses, clinically annotated cohorts, cellular compartments, and spatial tissue contexts. Rather than presenting each dataset as an independent resource, this organization enables systematic evaluation of whether molecular and pathway-level signals are maintained, altered, or resolved differently across complementary biological contexts.

### 2.2. PDAC BioPortal performs cross-cohort characterization of treatment-naïve and treatment-associated transcriptional programs

To characterize coordinated biological programs distinguishing PDAC tumors from normal pancreas and transcriptional programs associated with treatment, we performed complementary gene set enrichment analysis (GSEA) and over-representation analysis (ORA) across tumor-versus-normal and pre-versus post-SBRT transcriptomic cohorts. GSEA was performed using ranked differential-expression profiles to identify coordinated transcriptional changes, whereas ORA evaluated biological processes enriched among significantly differentially expressed genes.

Comparison of primary PDAC tumors (TCGA-PAAD) against normal pancreas (GTEx) revealed complementary pathway signatures across enrichment frameworks (Figure 2). Rank-sum enrichment analysis (ZGSEA; displayed as NES in the portal UI) was dominated by cell-cycle and proliferative programs, including mitotic metaphase/anaphase, cell-cycle checkpoints, E2F targets, and G2-M checkpoint, alongside neutrophil degranulation (all FDR < 1.71 × 10⁻⁵; Figure 2A-B). Leading-edge overlap analysis demonstrated shared core genes linking TNF-α/NF-κB, apoptosis, and DNA-repair signatures (Figure 2C-D). By contrast, ORA of significantly differentially expressed genes highlighted epithelial-mesenchymal transition (EMT), extracellular matrix (ECM) organization, inflammatory signaling, interferon responses, complement, and KRAS-associated programs (Figure 2E-G). These distinct emphases reflect the complementary mechanics of rank-based GSEA versus threshold-based ORA, with both frameworks converging on immune and inflammatory processes.

**Figure 2.**
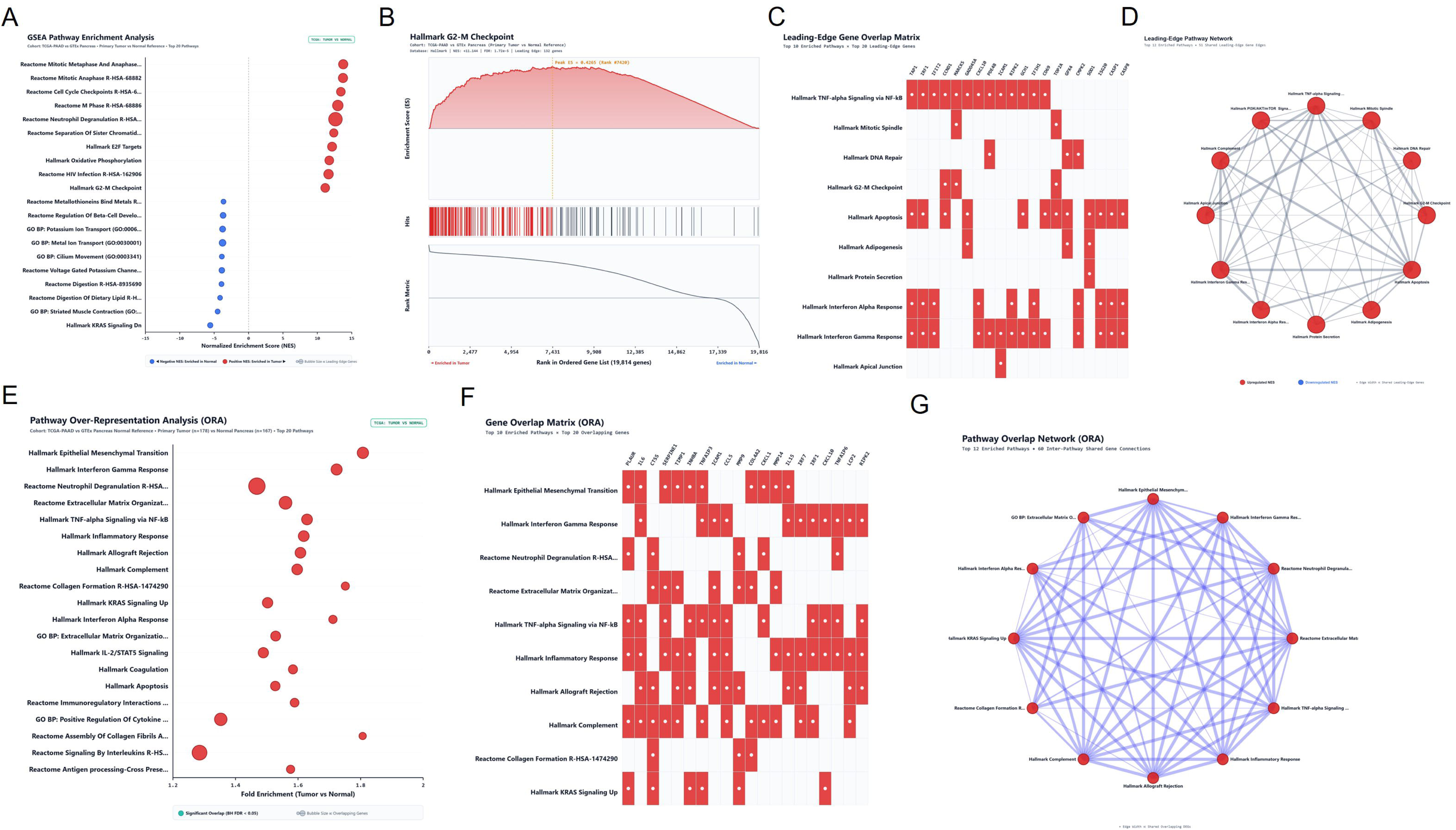
Pathway-level characterization of transcriptional alterations between primary PDAC tumors and normal pancreas. (A-D) Rank-sum pathway enrichment analysis (ZGSEA, displayed as NES in the portal UI) comparing TCGA primary PDAC tumors (n = 178) with GTEx normal pancreas specimens (n = 167). (A) Top enriched pathways across upregulated and downregulated programs. (B) Representative GSEA running enrichment plot for Hallmark G2-M Checkpoint (ZGSEA = +11.16 [portal UI NES], FDR < 1.71 × 10-5; 132 leading-edge genes; 1,000-permutation NESperm = +1.63, q < 0.001). (C) Leading-edge gene overlap matrix across top enriched Hallmark gene sets. (D) Pathway overlap network based on shared leading-edge genes across the top 12 enriched Hallmark pathways. (E-G) Complementary over-representation analysis (ORA) of significantly differentially expressed genes: (E) top 20 significantly over-represented pathways by fold enrichment, (F) gene overlap matrix of shared differentially expressed genes across top 10 pathways, and (G) ORA pathway overlap network connecting top 12 enriched pathways.

We next compared pre-treatment biopsies against post-neoadjuvant surgical specimens in the SBRT-containing cohort (GSE225767; Figure 3). Rank-sum pathway analysis identified prominent positive enrichment of mesenchymal and stromal remodeling programs, led by EMT, collagen biosynthesis, ECM organization, myogenesis, and IL-2/STAT5 signaling (all FDR < 1.31 × 10⁻⁴), alongside moderate enrichment of angiogenesis and hypoxia (Figure 3A-B). Conversely, G2-M checkpoint, interferon-alpha response, and heme metabolism were negatively enriched, reflecting higher expression in pre-treatment biopsies rather than active pathway inhibition. Leading-edge overlap and ORA independently confirmed tight gene-level connectivity across EMT, collagen, and ECM remodeling programs (Figure 3C-G).

**Figure 3.**
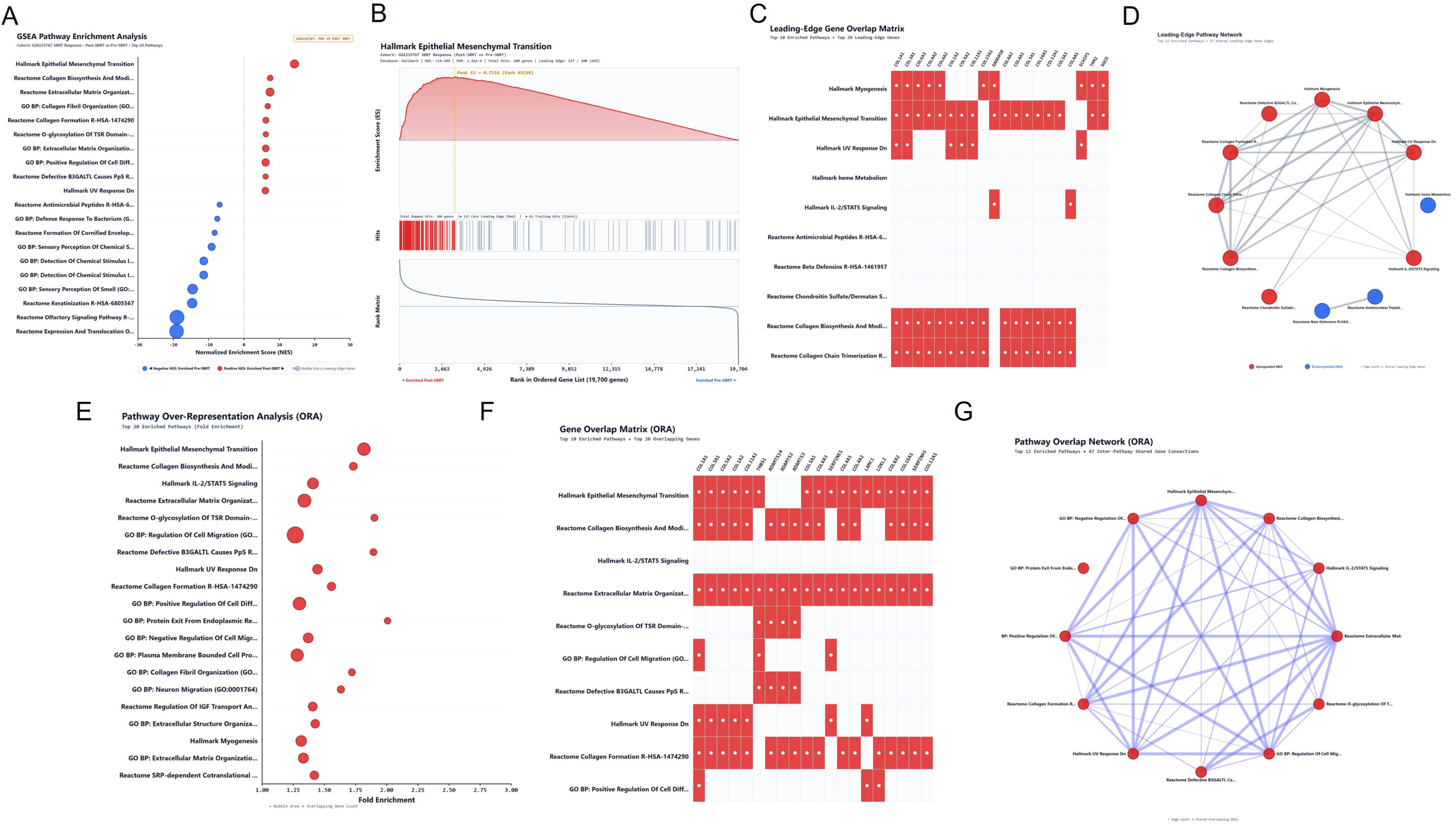
Pathway-level characterization of transcriptional responses associated with neoadjuvant SBRT in PDAC. (A-D) Rank-sum pathway enrichment analysis (ZGSEA, displayed as NES in the portal UI) comparing post-SBRT surgical resections (n = 29) with pre-SBRT FNA biopsies (n = 26) in GSE225767. (A) Top 20 positively and negatively enriched pathways ranked by rank-sum enrichment Z-score (ZGSEA, labeled NES in the portal UI). (B) Representative GSEA running enrichment plot for Hallmark Epithelial Mesenchymal Transition (ZGSEA = +14.29, FDR < 1.31 × 10-4; 137/200 leading-edge genes; 1,000-permutation NESperm = +1.94, q = 0.006). (C) Leading-edge gene overlap matrix across top 10 enriched Hallmark and Reactome gene sets. (D) Leading-edge pathway overlap network connecting top 12 enriched pathways. (E-G) Complementary ORA of significantly differentially expressed genes between pre-SBRT and post-SBRT specimens: (E) top 20 over-represented pathways by fold enrichment, (F) gene overlap matrix across top 10 enriched pathways, and (G) ORA pathway overlap network across top 12 pathways.

In support, ORA revealed enrichments for genes associated with EMT, collagen biosynthesis, extracellular matrix organization, collagen formation, cell migration, and other related extracellular remodeling annotations among significantly differentially expressed genes (Figure 3E-G). The common subset of genes underlying these annotations accentuated the cross-talk among the identified mesenchymal and extracellular matrix signatures.

The combined analysis of GSEA and ORA identifies a transcriptional signature associated with post-treatment bulk-tissue samples, marked by dominant mesenchymal and extracellular matrix related programs. While these signatures imply that large changes are happening between pre-and post-treatment samples, bulk RNA-seq cannot discriminate whether the signature here is mediated by changes at the level of cells (e.g., in expression levels) vs. changes in stromal / immune composition, or both. Finally, because treatment-associated comparisons are included in the cohort, these signatures do not demonstrate that SBRT exclusively drove the documented changes.

To compare functional architectures across disease and treatment contexts, we evaluated 1,000-permutation enrichment profiles (NESperm) across all 50 MSigDB Hallmark pathways (Supplementary Table S5). Primary PDAC tumorigenesis and neoadjuvant treatment remodeling exhibited a modest inverse rank correlation (Spearman ρ = −0.303, p = 0.032), sharing selective enrichment of EMT (leading-edge Jaccard index = 0.517) and angiogenesis. By contrast, independent treatment-associated cohorts (bulk SBRT GSE225767 and whole-sample single-nucleus chemoradiotherapy pseudobulk GSE202051; n = 14 CRT vs. n = 18 naïve) demonstrated significant global rank-order concordance (Spearman ρ = 0.512, p = 1.46 × 10⁻⁴; Pearson r = 0.195, p = 0.175). Compartment-resolved single-nucleus deconvolution showed that whole-sample negative shifts masked lineage-specific positive enrichments, such as EMT in CD8+ T cells and macrophages, TNF-α/inflammatory signaling in vascular endothelium, and myogenesis in pericytes, alongside negative cell-cycle enrichment in malignant epithelium (Supplementary Table S5). Patient-level composition analysis confirmed significant malignant epithelial depletion following CRT (49.6% to 14.5%; CLR FDR q = 0.0061), whereas apparent relative expansions in non-malignant lineages were non-significant under CLR transformation (CLR q = 0.271-0.880; Supplementary Table S2). Kitagawa-Oaxaca-Blinder decomposition further demonstrated that whole-sample cell-cycle suppression largely reflected parenchymal debulking (e.g., MYC targets v2: 96.4% compositional), whereas stress and immune shifts (such as reactive oxygen species: 99.1% within-cell) reflected cell-intrinsic transcriptional adaptation (Supplementary Table S5).

### 2.3. Analysis of NRF2-associated redox network as a biological case study using PDAC BioPortal

Given the established roles of oxidative stress and redox adaptation in PDAC biology and therapeutic resistance, together with our pathway analyses highlighting alterations in cell-cycle regulation and EMT, we selected the NRF2-associated transcriptional network for detailed interrogation using PDAC BioPortal. The integrated cross-cohort and cross-modal analyses provided a framework to investigate whether NRF2-associated redox programs exhibited consistent or context-dependent patterns across primary tumors, treatment-associated transcriptional changes, and distinct cellular compartments. This analysis also allowed us to examine how NRF2-associated programs related to broader metabolic and stress-response adaptations in PDAC.

#### 2.3.1. The NRF2-associated transcriptional network shows broad transcriptional elevation in primary PDAC

Having established an interactive framework for custom gene-set interrogation, we used PDAC BioPortal to investigate the transcriptional landscape of NRF2-associated redox regulation in primary PDAC by examining a curated 20-gene signature comprising five functionally related components: core NRF2 regulatory machinery, canonical antioxidant response genes, glutathione synthesis and transport, NADPH-generating metabolism, and iron homeostasis and ferroptosis-associated defense (Figure 4A). The gene panel was assembled based on previously reported NRF2-associated regulatory relationships and established roles in cellular antioxidant defense and redox homeostasis ^[24–26]^.

**Figure 4.**
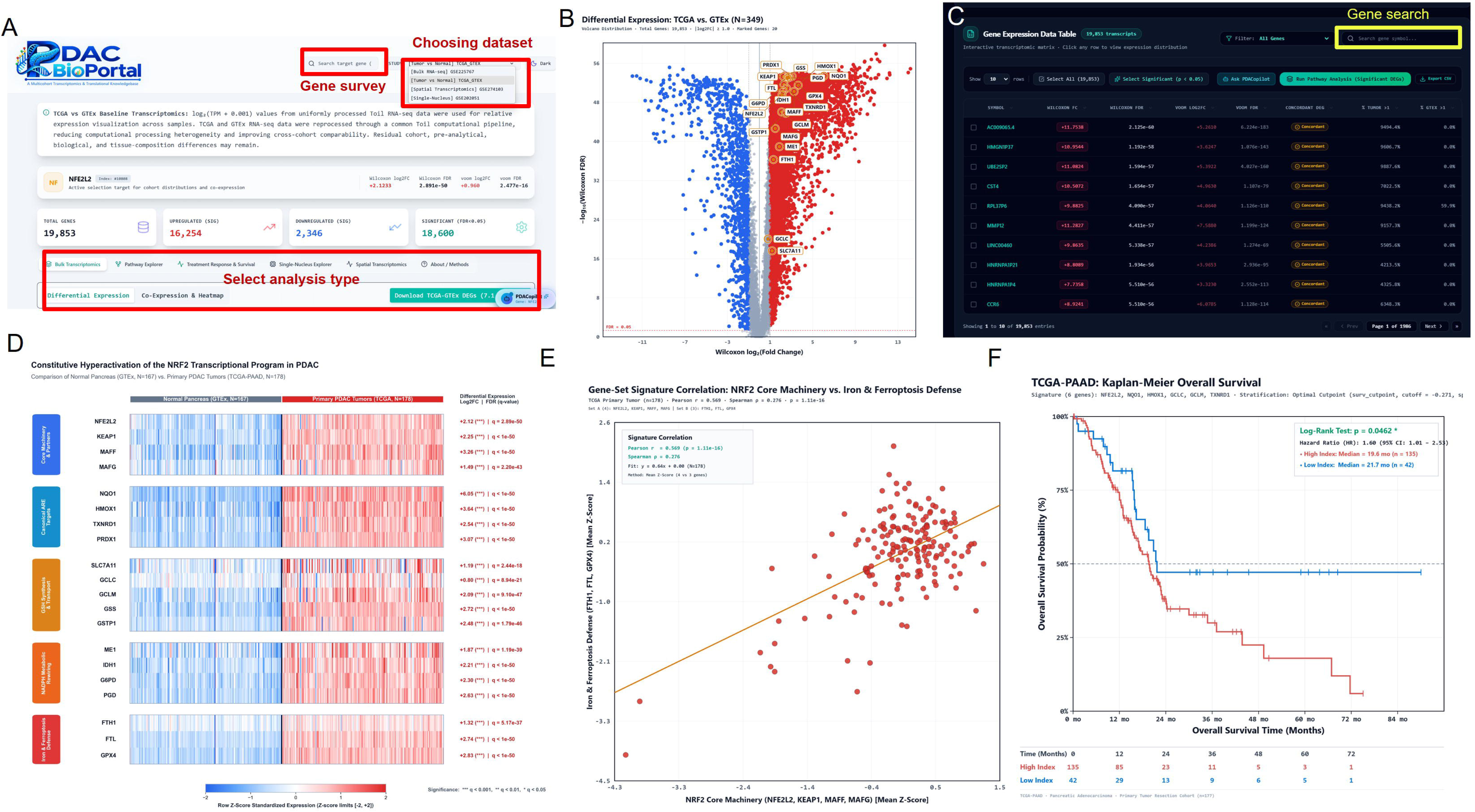
PDAC BioPortal interrogation of the NRF2-associated redox network in primary PDAC. (A) Interface displaying TCGA-GTEx baseline cohort selection for NFE2L2. (B) Volcano plot comparing TCGA primary PDAC (n = 178) vs. GTEx normal pancreas (n = 167; 19,853 genes) highlighting the 20 curated NRF2-panel genes (all FDR q < 10⁻¹⁷; Table S1). (C) Interactive differential-expression table with Wilcoxon and limma-voom metrics. (D) Row Z-score heatmap of the 20-gene program across normal (n = 167) and tumor (n = 178) samples. (E) Correlation between NRF2 Core Machinery and Iron/Ferroptosis Defense signatures (n = 178; r = 0.569, p = 1.11 × 10⁻¹⁶). (F) Exploratory Kaplan-Meier curve in TCGA-PAAD (N = 177) for the six-gene NRF2 signature (unadjusted log-rank p = 0.0462, maxstat padj = 0.393; primary continuous Cox HR = 1.35, p = 0.0422).

Comparison of primary PDAC tumors (n = 178) with normal pancreas (n = 167) revealed coordinated, transcriptome-wide elevation across all 20 genes of the NRF2-associated panel (all Wilcoxon FDR q < 10⁻¹⁷; Figure 4B-D; complete statistics in Supplementary Table S1). Core regulatory components (NFE2L2, KEAP1, MAFF, and MAFG) and canonical antioxidant targets, led by NQO1, HMOX1, PRDX1, and TXNRD1, were uniformly upregulated in primary tumors ^[24,26].^

Across the remaining functional modules (Figure 4D; Supplementary Table S1), glutathione synthesis and cystine-transport genes (GSS, GSTP1, GCLM, SLC7A11, and GCLC), NADPH-generating enzymes (ME1, IDH1, G6PD, and PGD), and iron-homeostasis/ferroptosis-defense genes (FTH1, FTL, and GPX4) were consistently elevated in primary PDAC.

Gene-signature correlation analysis revealed a positive association between the four-gene NRF2 core machinery and three-gene iron/ferroptosis-defense modules in primary PDAC (Pearson r = 0.569, p = 1.11 × 10⁻¹⁶; Spearman ρ = 0.276; n = 178; Figure 4E; Supplementary Table S6), with the difference between linear and rank correlations reflecting right-skewed high-expressing tumors.

To explore potential clinical associations, we evaluated a six-gene NRF2 antioxidant signature (NFE2L2, NQO1, HMOX1, GCLC, GCLM, and TXNRD1) in TCGA-PAAD (N = 177; Supplementary Table S6). Modeled continuously in univariate Cox regression, higher signature expression was nominally associated with shorter overall survival (HR = 1.35 per unit Z-score, 95% CI: 1.01-1.81, Wald p = 0.0422; Schoenfeld PH test p = 0.690). Exploratory optimal-cutpoint Kaplan-Meier stratification showed a consistent unadjusted separation (Figure 4F), although this did not survive multiple-cutpoint adjustment (maxstat padj = 0.393) and multivariable staging adjustment was precluded by incomplete clinical annotations. By contrast, the three-gene iron/ferroptosis-defense signature (FTH1, FTL, GPX4) showed no significant survival association continuously (Cox HR = 1.07, p = 0.630, PH p = 0.872) or by cutpoint stratification (maxstat padj = 0.590; Supplementary Figure S1A; Supplementary Table S6). These survival results therefore represent exploratory transcriptional associations rather than validated prognostic biomarkers.

#### 2.3.2. NRF2-associated programs exhibit context-dependent remodeling following SBRT

We next used the SBRT module of PDAC BioPortal to examine whether the NRF2-associated redox transcriptional program was altered following multimodal neoadjuvant treatment incorporating SBRT. Genome-wide differential-expression analysis of the pre-vs post-SBRT cohort revealed substantial transcriptional differences between pre-SBRT (n = 26) and post-SBRT (n = 29) tumors (Figure 5A-C). However, interrogation of the curated NRF2-associated gene panel showed that selected components of the NRF2-associated transcriptional program exhibit treatment-associated changes rather than a uniform shift (Figure 5E).

**Figure 5.**
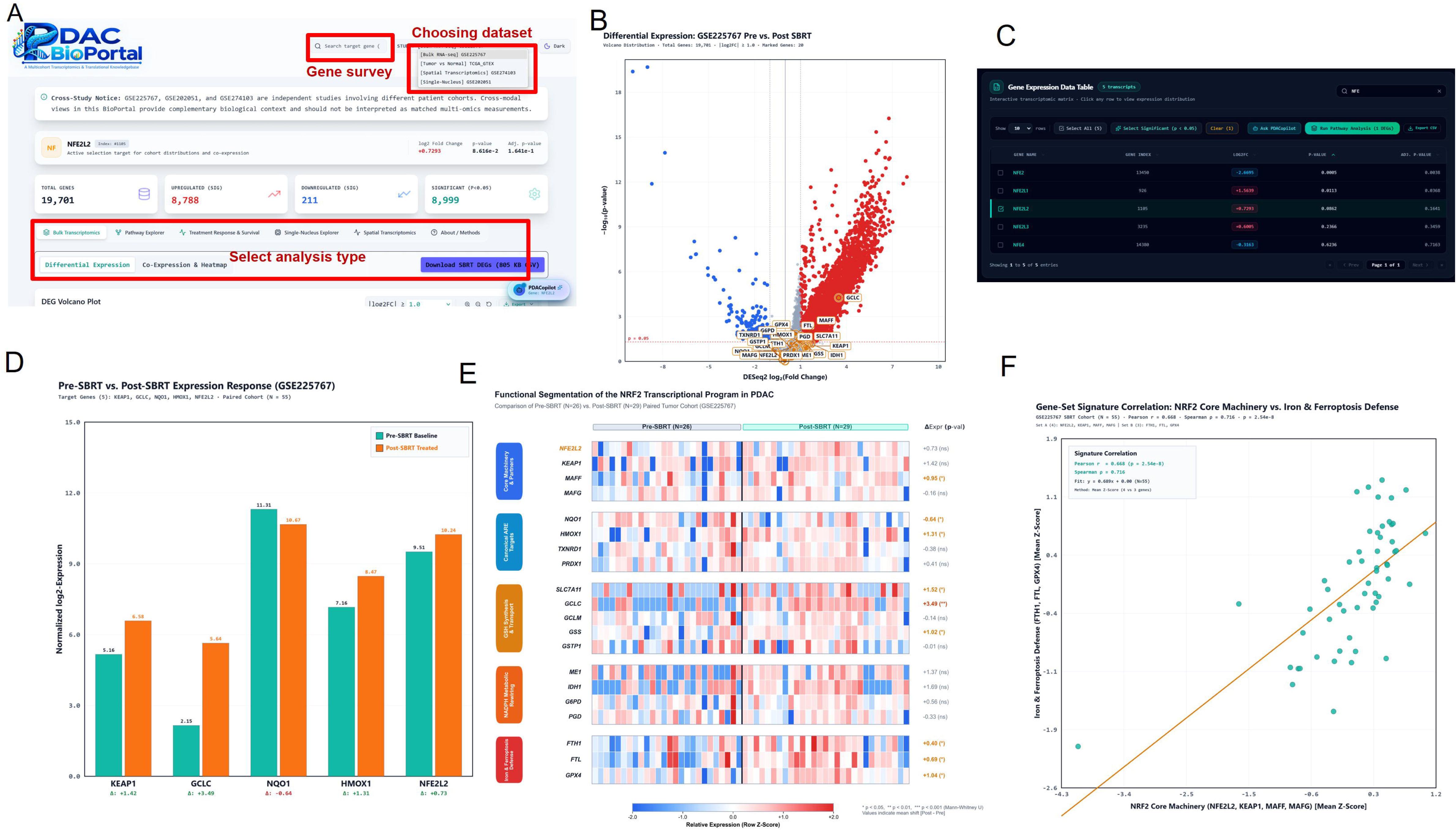
Interrogation of NRF2-associated remodeling in the GSE225767 neoadjuvant cohort. (A) Portal SBRT interface for NFE2L2. (B) DESeq2 volcano plot comparing post-neoadjuvant resections (n = 29) vs. pre-treatment FNA biopsies (n = 26; 19,701 genes). (C) Interactive DESeq2 statistics table. (D) Mean log2 expression shifts (Δ) for representative genes. (E) Row Z-score heatmap of the 20-gene NRF2 panel annotated with nominal Mann-Whitney p-values (*p < 0.05, **p < 0.01; GCLC passes 20-gene BH q = 0.035; Table S1). (F) Signature correlation between NRF2 Core Machinery and Iron/Ferroptosis Defense (N = 55; r = 0.668, p = 2.54 × 10⁻⁸).

In contrast to the uniform upregulation seen in primary PDAC, the 20-gene NRF2 panel underwent selective, heterogeneous remodeling between pre-treatment biopsies (n = 26) and post-neoadjuvant surgical specimens (n = 29) in GSE225767 (Figure 5B-E; complete statistics in Supplementary Table S1). Neither the core regulatory machinery (NFE2L2, KEAP1, MAFF, MAFG) nor canonical antioxidant targets (HMOX1, NQO1, TXNRD1, PRDX1) or NADPH-generating enzymes (ME1, IDH1, G6PD, PGD) passed panel-wide FDR correction, with HMOX1 and NQO1 showing divergent nominal trends.

Among glutathione and iron/ferroptosis-defense components, FDR-significant post-treatment elevation was restricted to GCLC (20-gene BH q = 0.035; DESeq2 adj. p = 6.64 × 10⁻⁴) and FTL in genome-wide DESeq2 testing (adj. p = 0.044). By contrast, SLC7A11, GSS, FTH1, and GPX4 showed only nominal upward shifts that did not survive panel-wide FDR correction, while GCLM and GSTP1 remained unchanged (Figure 5D-E; Supplementary Table S1).

Despite heterogeneous single-gene shifts in GSE225767, the positive correlation between NRF2 core machinery and iron/ferroptosis-defense signatures persisted across the treatment-associated cohort (Pearson r = 0.668, Spearman ρ = 0.716, p = 2.54 × 10⁻⁸; N = 55; Figure 5F; Supplementary Table S6).

At the pathway level, Reactome KEAP1-NFE2L2 signaling was strongly enriched in primary PDAC versus normal pancreas (ZGSEA = +8.40, FDR < 1.71 × 10⁻⁵; Supplementary Figure S1B), whereas GO cellular response to oxidative stress showed only nominal post-treatment enrichment in GSE225767 that did not pass FDR correction (ZGSEA = +2.07, nominal p = 0.0195, FDR = 0.131; Supplementary Figure S1C; Supplementary Table S6).

Finally, an eight-gene NRF2 target signature correlated strongly with the 10-gene Radiation Sensitivity Index (RSI) ^[27]^ in both primary PDAC (r = 0.869, p = 1.15 × 10⁻⁵⁵) and GSE225767 (r = 0.743, p = 8.28 × 10⁻¹¹; Supplementary Figure S1D-E; Supplementary Table S6), establishing transcriptional covariance rather than causal radioresistance.

#### 2.3.3. Cellular localization of NFE2L2 expression reveals compartment-specific remodeling in PDAC

To determine the cellular contexts underlying the heterogeneous NFE2L2-associated transcriptional patterns identified across the bulk transcriptomic cohorts, we next leveraged the single-nucleus analysis module of PDAC BioPortal to examine NFE2L2 expression at cellular resolution in the single-nucleus neoadjuvant-treated PDAC cohort (Figure 6). This framework enabled interrogation of NFE2L2 expression across major cellular populations and annotated sub-lineages, as well as comparisons across treatment contexts and patient-level pseudobulk profiles.

**Figure 6.**
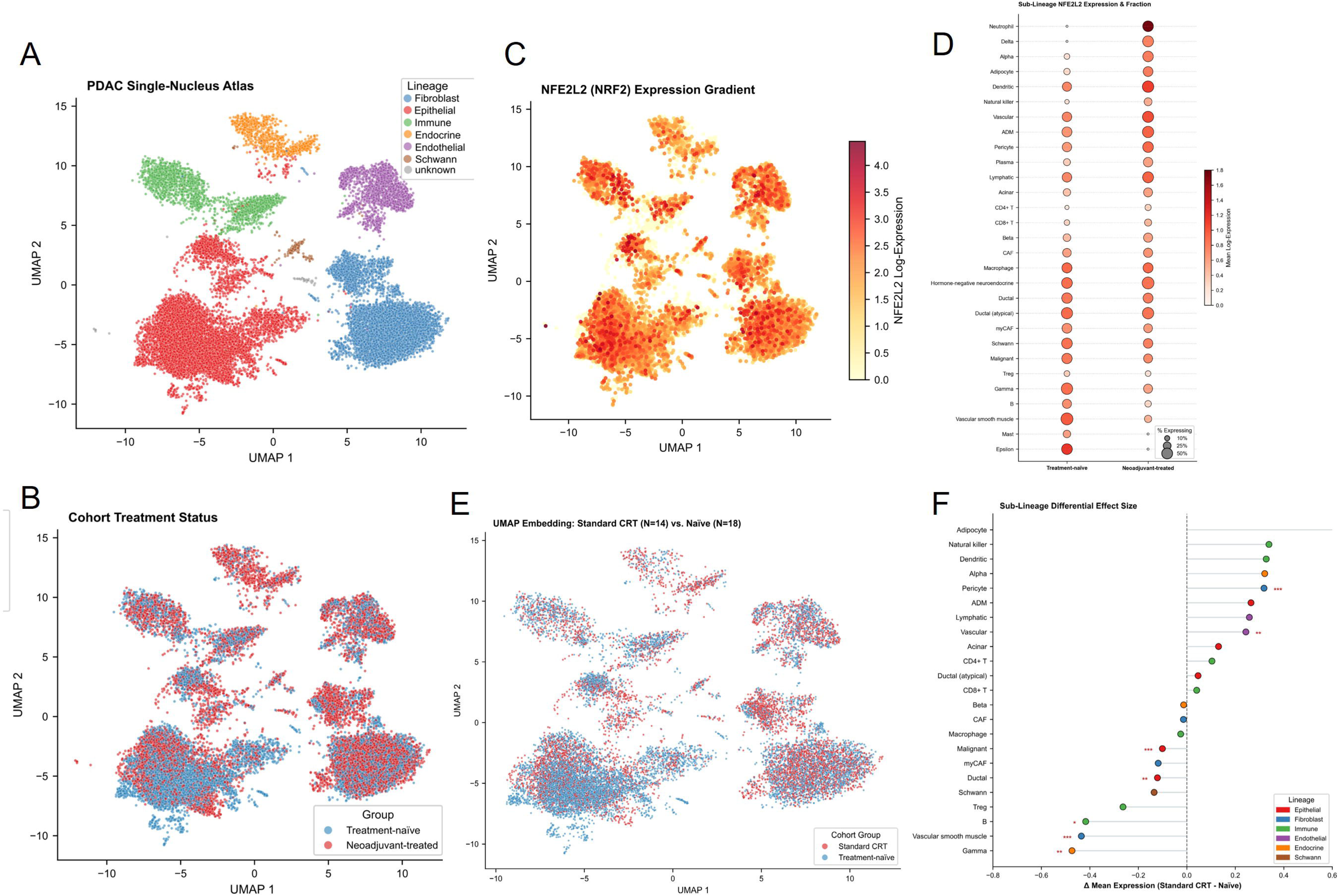
Single-nucleus analysis reveals cellular and treatment-associated patterns of NFE2L2 expression in PDAC (GSE202051). (A) UMAP visualization of the PDAC single-nucleus atlas colored by major cellular lineage. (B) UMAP embedding colored by cohort treatment status (treatment-naïve vs. neoadjuvant-treated). (C) UMAP feature plot of NFE2L2 log-normalized expression across cellular compartments. (D) Bubble plot comparing mean NFE2L2 log-expression (color) and fraction of expressing nuclei (bubble size) across 29 annotated cellular sub-lineages in treatment-naïve versus neoadjuvant-treated tumors. (E) UMAP projection comparing nuclei from patients receiving standard chemoradiotherapy (CRT; n = 14) versus treatment-naïve patients (n = 18). (F) Sub-lineage differential expression analysis (standard CRT minus treatment-naïve mean expression) colored by cellular lineage, with significance asterisks indicating statistically supported differences.

Across the GSE202051 atlas, neoadjuvant-treated specimens exhibited shifted cellular proportions toward endothelial, fibroblast, and endocrine lineages alongside compartment-dependent NFE2L2 expression (Figure 6A-D). At patient-level pseudobulk resolution comparing all neoadjuvant-treated (n = 25) versus treatment-naïve (n = 18) tumors, NFE2L2 increased significantly in dendritic, delta, vascular endothelial, and alpha sub-lineages (FDR q = 0.027-0.047), while showing a nominal decrease in malignant epithelial cells (p = 0.02; Supplementary Figure S2A; Supplementary Table S6).

Restricting the comparison to the homogeneous standard CRT subgroup (n = 14) versus treatment-naïve tumors (n = 18) revealed reduced NFE2L2 expression across malignant epithelial, ductal, B cell, gamma islet, and vascular smooth muscle populations (with the fraction of NFE2L2-positive malignant nuclei declining from ∼45% to ∼35%), alongside significant increases in pericytes and vascular endothelial cells (Figure 6E-F; Supplementary Figure S2B; Supplementary Table S6).

#### 2.3.4. Spatial transcriptomic analysis reveals heterogeneous distribution of NFE2L2 expression across treatment-naïve PDAC tissues

To further understand the expression distribution of the NRF2-associated transcriptional program into the spatial context, we used the spatial transcriptomics module of PDAC BioPortal to interrogate NFE2L2 expression across five treatment-naïve PDAC specimens from the GSE274103 dataset (Figure 7). The portal enabled sequential selection of individual datasets, patients, tissue sections, and genes, followed by visualization of expression directly within the corresponding tissue architecture (Figure 7A-B).

**Figure 7.**
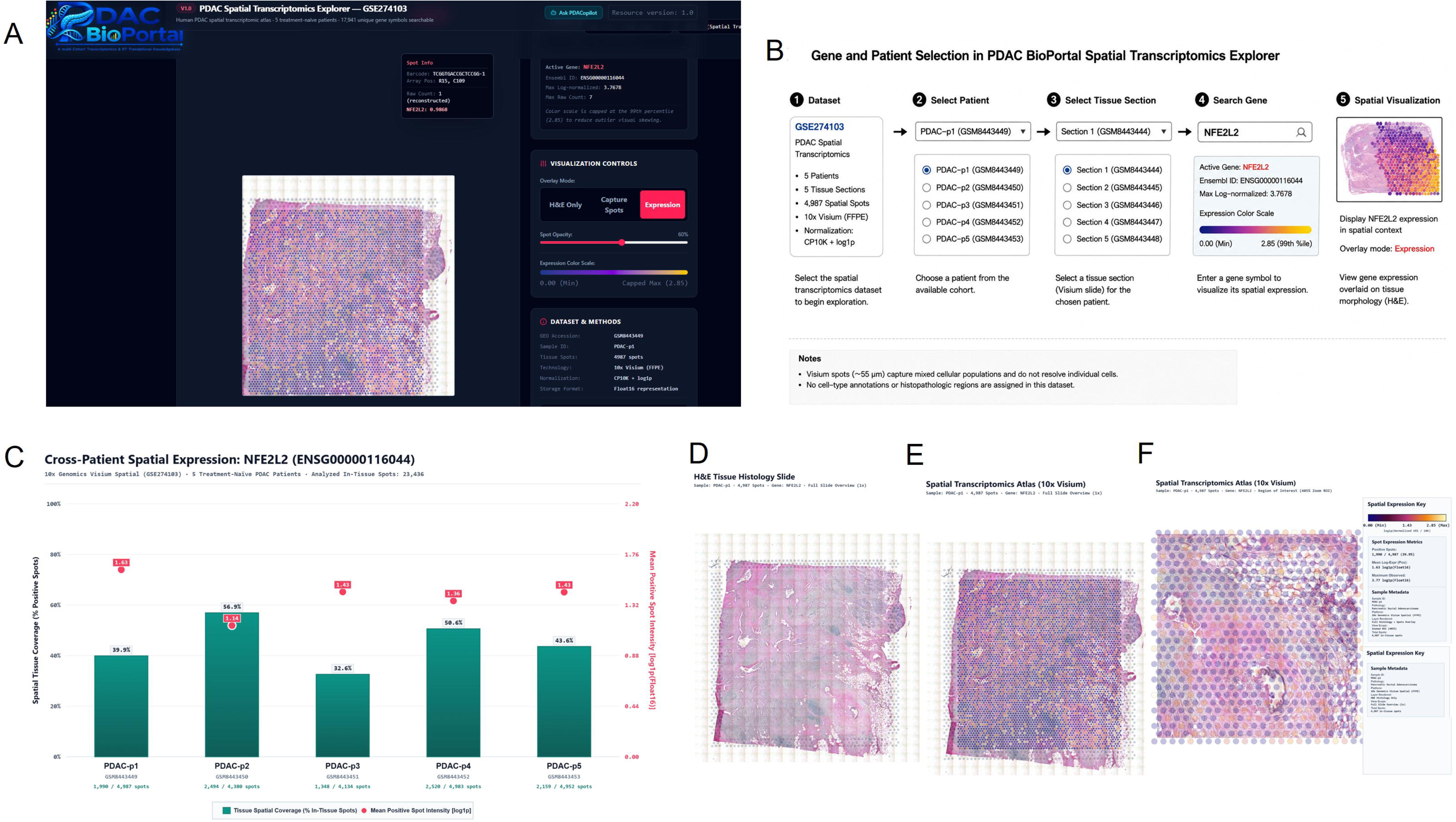
Spatial transcriptomic analysis of NFE2L2 expression across treatment-naïve PDAC tissues (GSE274103). (A) PDAC BioPortal Spatial Transcriptomics Explorer interface. (B) Sequential selection workflow from dataset, patient, and tissue section to gene query and spot-level visualization. (C) Cross-patient summary of NFE2L2 spatial tissue coverage (% positive spots, bars) and mean positive-spot intensity (log1p Float16, red circles) across five treatment-naïve PDAC sections (PDAC-p1 to PDAC-p5; 23,436 total spots). (D) Brightfield H&E histology overview of representative specimen PDAC-p1 (GSM8443449; 4,987 spots). (E) Full-slide spatial overlay of NFE2L2 spot-level expression on PDAC-p1. (F) Enlarged region of interest (405% zoom) illustrating focal spatial heterogeneity across Visium capture spots.

Across the five treatment-naïve PDAC sections, NFE2L2 transcripts were detected in 10,511 of 23,436 in-tissue Visium spots (44.8% overall coverage; Figure 7C; per-patient metrics in Supplementary Table S3). Spatial coverage varied substantially across tumors, ranging from 32.6% (PDAC-p3) to 56.9% (PDAC-p2), whereas mean positive-spot expression intensity remained within a narrower range (1.142-1.627 log1p CP10k; maximum spot intensity 3.146-4.234). Notably, PDAC-p2 combined the widest spatial coverage with the lowest mean positive-spot intensity, whereas PDAC-p1 showed the highest mean intensity over a more focal distribution.

Together, these spatial measurements distinguish the proportion of tissue regions expressing NFE2L2 from local transcript abundance within positive spots. Because this module analyzes treatment-naïve specimens, it establishes a baseline spatial reference rather than evidence of treatment-induced spatial remodeling.

Overlaying NFE2L2 expression onto brightfield H&E sections confirmed focal spatial heterogeneity across all five tumors (Figure 7D-F). Because 10x Visium capture spots (55 μm) aggregate transcripts from multiple neighboring cells, these profiles establish tissue-region heterogeneity rather than cell-type-specific origin.

### 2.4. AI-assisted exploration with PDACopilot

To further improve accessibility and search functions, while facilitating biological interpretation of multimodal transcriptomic data, PDAC BioPortal incorporates PDACopilot, an integrated artificial intelligence (AI)-assisted research assistant that enables conversational exploration of transcriptomic datasets through natural-language queries. Unlike conventional visualization tools that require users to manually navigate multiple analytical modules, PDACopilot provides context-aware interpretation of transcriptomic results, summarizes biological observations, and guides users toward complementary analyses across the integrated datasets/modules. This AI-assisted workflow lowers the computational barrier for exploratory analysis while maintaining direct linkage to the underlying transcriptomic evidence (Supplementary Figure S3).

#### 2.4.1. Deterministic Architecture and Data-Grounded Querying

To democratize multi-modal transcriptomic exploration without requiring programmatic bioinformatics scripting, we developed PDACopilot, an autonomous, data-grounded AI copilot embedded directly within the PDAC BioPortal architecture. Unlike generic large language models (LLMs) that are vulnerable to numerical hallucinations, statistical sign reversals, and fabricated literature citations, PDACopilot operates under a deterministic, evidence-grounded retrieval pipeline (Supplementary Figure S3): Intent and Cohort Routing: Natural language prompts from investigators are parsed by an IntentRouter that classifies queries into discrete analytical tasks (e.g., tumor vs normal comparison, radiotherapy treatment response, cell type lineage expression, spatial localization, and cross module synthesis).

Programmatic QueryEngine Execution: Rather than generating figures or metrics from memory, PDACopilot calls deterministic TypeScript/WebAssembly tools (QueryEngine.ts) that query the portal’s pre-computed differential expression tables, Float16/Float32 binary matrices, and spatial coordinate stores with sub-15 millisecond latency.

Evidence Consistency Validation: Retrieved empirical metrics (Wilcoxon and limma-voom log2FC, FDR q-values, group means, single-nucleus lineage fractions, and Visium spot metrics) are compiled into structured evidence objects (EvidenceValidator.ts) that constrain generated responses (Supplementary Table S4). On a frozen 20-question diagnostic benchmark, evidence locking improved consistency from unconstrained Gemini 2.5 Flash (20.0/40, 50.0%; exhibiting FDR inversions, question drift, an unpaired-cohort error, and a fabricated table) and Llama 3.3 70B (15.2/40, 38.1%) to 39.0/40 (97.5%; 0 numerical hallucinations and 0 statistical errors), and achieved 342/350 (97.7%) on an expanded 35-question, 10-criterion analytical suite.

#### 2.4.2. Cross-modal interrogation of NFE2L2 expression across treatment contexts

To demonstrate cross-modal exploration using PDACopilot, NFE2L2 was selected as a representative gene for interrogation across the integrated PDAC datasets. The retrieved evidence highlighted distinct expression patterns across biological contexts. NFE2L2 was significantly elevated in primary PDAC compared with normal pancreas, whereas its increase in post-treatment bulk specimens was not statistically significant. At the single-nucleus level, sub-lineage and patient-level pseudobulk analyses identified significant increases in NFE2L2 expression within vascular endothelial, pericyte, dendritic, and islet endocrine subsets in the neoadjuvant-treated cohort, whereas malignant epithelial expression was reduced. Spatial transcriptomic analysis further detected NFE2L2 transcripts across all five treatment-naïve specimens, with variation in maximum spot-level expression. Together, these observations illustrate how PDACopilot can retrieve and connect gene-level evidence across independent bulk, single-nucleus, and spatial datasets while preserving the statistical and experimental context of each analysis. The findings demonstrate context-dependent NFE2L2 expression patterns but do not establish cell-specific NRF2 protein activity, a radiation-specific response, or a causal relationship between endothelial or stromal expression and treatment resistance.

### 2.5. Comparison with Existing Cancer Transcriptomic Resources

To contextualize PDAC BioPortal, we benchmarked its native capabilities against GEPIA2, cBioPortal, TISCH2, and CZ CELLxGENE (Table 3). While these pan-cancer resources excel within individual modalities, including TCGA/GTEx bulk queries (GEPIA2), multidimensional tumor genomics (cBioPortal), and pan-tissue single-cell visualization (TISCH2, CELLxGENE), none integrates baseline tumor-normal expression, neoadjuvant SBRT/CRT cohorts, patient-level pseudobulk deconvolution, interactive GSEA/ORA, survival modeling, and Visium spatial mapping within a unified PDAC workflow.

**Table 3.** Comparison of native capabilities between PDAC BioPortal and existing transcriptomic resources (assessed March 2026).

| **Capability / analytical feature** | **PDAC BioPortal** | **GEPIA2** | **cBioPortal** | **TISCH2** | **CELLxGENE** |
|---|---|---|---|---|---|
| Dedicated PDAC focus | ✓ | - | - | △ | △ |
| Harmonized PDAC tumor-versus-normal bulk RNA-seq | ✓ | ✓ | △ | - | - |
| PDAC neoadjuvant treatment-associated bulk transcriptomics | ✓ | - | △ | - | - |
| Pre-/post-treatment SBRT cohort interrogation | ✓ | - | - | - | - |
| PDAC single-nucleus reference atlas | ✓ | - | - | ✓ | ✓ |
| Treatment-associated single-nucleus analysis | ✓ | - | - | △ | △ |
| Patient-level pseudobulk differential expression | ✓ | - | - | △ | △ |
| Interactive spatial transcriptomic visualization | ✓ | - | - | - | △ |
| Interactive pathway enrichment (GSEA/ORA) | ✓ | △ | △ | △ | △ |
| Gene-level expression correlation | ✓ | △ | ✓ | △ | △ |
| Gene-set / signature correlation | ✓ | - | - | - | - |
| Gene-expression-based survival analysis | ✓ | ✓ | ✓ | - | - |
| Custom gene-set / signature survival analysis | ✓ | - | △ | - | - |
| Responder-versus-non-responder expression analysis | ✓ | - | △ | - | - |
| Integrated cross-modal gene navigation | ✓ | - | - | - | △ |
| Zero-programming web-based graphical interface | ✓ | ✓ | ✓ | ✓ | ✓ |
| AI-assisted natural-language exploration (PDACopilot) | ✓ | - | - | - | - |
| Evidence-grounded cross-modal result synthesis | ✓ | - | - | - | - |

✓: Fully supported as a native, dedicated interactive feature.
△: Limited, indirect, or pan-cancer unstructured support.
-: Not supported.

By pairing these multi-scale analytical layers with deterministic browser-side execution, reproducible exports, and PDACopilot’s evidence-grounded natural-language querying, PDAC BioPortal lowers technical barriers to cross-modal hypothesis generation without replacing external repositories.

PDAC BioPortal also incorporates PDACopilot, a data-grounded conversational interface designed to facilitate natural-language exploration of the portal’s integrated datasets. PDACopilot connects language-based queries with a deterministic bioinformatics QueryEngine, allowing users to retrieve underlying quantitative results, interrogate pathway-level associations, and synthesize observations across analytical modalities. Together with browser-based analysis, reproducible statistical outputs, and figure export functionality, these features extend the accessibility of multimodal PDAC transcriptomic analysis.

## 3. Discussion

A central barrier in translational pancreatic cancer research is that neoadjuvant therapy fundamentally restructures the tumor ecosystem rather than merely shrinking or intensifying the untreated tumor state. In many biomarker studies, post-treatment bulk RNA sequencing is interpreted as though tissue-level fold changes reflect uniform reprogramming inside surviving cancer cells. By connecting primary tumors, neoadjuvant bulk cohorts, single-nucleus profiles, and spatial tissue architecture within PDAC BioPortal, our work demonstrates why that assumption breaks down in PDAC: physical clearance of malignant epithelium and accompanying fibroinflammatory remodeling dominate the bulk transcriptome, causing tumor-cell adaptations and microenvironmental turnover to be misattributed unless evaluated across biological scales.

The conservation of pathway hierarchies across distinct neoadjuvant regimens (ablative SBRT and conventional chemoradiotherapy) carries direct translational weight for radiation oncology ^[5,7,8,10]^. Rather than representing protocol-specific noise, epithelial plasticity, extracellular matrix remodeling, and microvascular inflammatory activation emerge as shared biological consequences of cytotoxic and radiation stress in human PDAC. Crucially, separating compositional turnover from within-cell signaling distinguishes two therapeutic problems that bulk profiling routinely blurs together ^[4,17]^. First, apparent suppression of proliferative programs after neoadjuvant therapy primarily reflects parenchymal debulking rather than durable cell-cycle arrest inside surviving tumor clones, warning against using bulk proliferation decline alone as a surrogate for eradication of resistant residual disease. Second, persistent oxidative-stress, inflammatory, and endothelial activation programs represent genuine cell-intrinsic stress adaptation within surviving microenvironmental niches, pointing toward stroma-directed, vascular-modulating, or immune-combinatorial strategies as rational adjuncts to neoadjuvant radiotherapy.

Our NRF2 redox case study demonstrates how this multi-scale perspective sharpens therapeutic hypotheses rather than simply cataloging differential expression ^[24–26]^. In treatment-naïve PDAC, coordinated elevation of cystine uptake, glutathione biosynthesis, NADPH regeneration, and GPX4-dependent lipid peroxide detoxification reflects a broad oncogene-driven antioxidant shield that buffers proliferating tumor cells against oxidative injury and ferroptosis ^[28–30]^. After neoadjuvant therapy, however, this global program narrows into a selective metabolic bottleneck centered on glutathione synthesis and iron handling alongside prominent vascular and myeloid redox activation. Biologically, this suggests two actionable research directions. First, rather than attempting broad upstream NRF2 blockade, which faces the challenge that NFE2L2 transcription declines in residual malignant epithelium after chemoradiotherapy and NRF2 activity is controlled primarily through KEAP1-mediated protein stabilization, radiosensitization efforts may gain greater leverage by targeting downstream metabolic vulnerabilities such as glutamate-cysteine ligase (GCLC), cystine transport, or iron-dependent ferroptosis induction ^[31]^. Second, heightened vascular endothelial and perivascular redox signaling in treated tumors raises the testable hypothesis that microvascular antioxidant buffering preserves endothelial survival during radiation, motivating mechanistic studies into whether redox-targeted interventions reshape tumor perfusion and immune infiltration after SBRT.

Beyond these specific biological leads, the practical payoff of PDAC BioPortal is disciplined target triage prior to costly experimental investment. Before launching organoid perturbations, murine radiation models, or clinical tissue staining, investigators can verify in seconds whether a candidate gene reflects malignant epithelial biology, stromal or endothelial remodeling, or focal spatial heterogeneity, while PDACopilot accelerates hypothesis exploration through evidence-grounded retrieval rather than unconstrained language generation ^[22,23,32–34]^. Important boundaries remain: the integrated datasets stem from independent clinical cohorts, the SBRT-containing cohort compares unpaired pre-treatment biopsies with post-treatment surgical resections exposed to combined systemic chemotherapy and radiation, the spatial module is limited to treatment-naïve tumors, and transcriptional associations cannot prove protein activity or causal resistance ^[18,19]^. Because the platform uses a modular static architecture, newly generated matched longitudinal cohorts, post-treatment spatial atlases, proteomics, and functional radiation perturbation screens can be incorporated seamlessly as they emerge, progressively moving pancreatic cancer bioinformatics from descriptive cataloging toward mechanism-guided radiosensitization.

In summary, PDAC BioPortal provides an open-access, multi-scale resource that separates tissue-composition artifacts from cell-intrinsic adaptation in pancreatic cancer, equipping researchers with a rigorous foundation for context-aware target prioritization and translational radiotherapy hypotheses.

## 4. Materials and Methods

### 4.1. System architecture and implementation

PDAC BioPortal was designed to enable multiscale exploration of candidate genes and transcriptional programs across complementary biological contexts in pancreatic ductal adenocarcinoma. The integrated datasets capture distinct levels of PDAC biology: bulk tumor and non-malignant pancreatic transcriptomes characterize broad disease-associated alterations; neoadjuvant treatment-associated cohorts enable investigation of therapy-induced remodeling;single-nucleus RNA sequencing (snRNA-seq) provides cellular-resolution context; and spatial transcriptomics resolves gene expression within native tissue architecture.

The current implementation links four analytical modules (baseline TCGA-GTEx, neoadjuvant SBRT-containing bulk GSE225767, single-nucleus GSE202051, and Visium spatial transcriptomics GSE274103) through a shared gene-centric interface (Table 1). Because these resources derive from independent clinical cohorts and platforms, cross-modal comparisons provide biological context and programmatic reproducibility rather than within-patient mechanistic equivalence. Browser implementation details are described in Section 4.7.

### 4.2. Data Sources and Data Preprocessing

Publicly available transcriptomic datasets were obtained from the UCSC Xena data repository and the NCBI Gene Expression Omnibus (GEO). Each dataset underwent standardized, dataset-specific preprocessing to harmonize gene nomenclature and ensure analytical consistency while preserving original sample metadata. Expression matrices were subsequently converted into web-optimized, indexed binary formats to enable sub-second browser-based visualization and client-side statistical querying within PDAC BioPortal.

#### 4.2.1. Tumor and non-malignant pancreatic transcriptomes

Harmonized RNA-sequencing data from pancreatic tumors and non-malignant pancreatic tissues were obtained from the UCSC Xena platform ^[35]^ via the TCGA-TARGET-GTEx Toil Recompute resource ^[36]^. The Toil processing pipeline processes raw sequencing reads through a unified computational framework, including STAR alignment and RSEM quantification, to eliminate technical batch effects and pipeline artifacts arising from disparate historical processing pipelines.

The portal cohort comprises 349 curated samples: 178 untreated primary TCGA-PAAD tumors ^[9]^, 167 GTEx normal pancreas specimens ^[20]^, and 4 TCGA solid-tissue normal controls (excluding one metastatic specimen). Baseline tumor-versus-normal differential expression evaluates the 178 primary tumors against the 167 GTEx normal samples (n = 345). Filtering 20,832 protein-coding genes for unmapped and zero-variance transcripts yielded 19,853 analyzed genes expressed as log2(TPM+0.001). Because TCGA and GTEx derive from distinct tissue-acquisition protocols, comparisons reflect broad disease-associated profiles rather than tumor-cell-intrinsic causation.

#### 4.2.2. Neoadjuvant treatment-associated PDAC transcriptomes

Bulk RNA-sequencing data capturing multimodal neoadjuvant treatment response incorporating stereotactic body radiotherapy were obtained from GEO accession GSE225767 ^[7]^. This study investigated transcriptional patterns associated with therapeutic response in patients with borderline resectable PDAC undergoing neoadjuvant multimodal therapy. Under the clinical trial protocol, patients received induction systemic chemotherapy followed by restaging and stereotactic body radiotherapy (SBRT; 30-33.6 Gy in 5 fractions) prior to pancreaticoduodenectomy and subsequent adjuvant chemotherapy. This cohort was therefore incorporated into PDAC BioPortal as a clinically annotated neoadjuvant treatment-associated transcriptomic resource rather than as an isolated in vivo model of radiation exposure alone.

The dataset comprises 55 unpaired bulk profiles: 26 pre-treatment fine-needle aspiration (FNA) biopsies and 29 post-treatment surgical resections from independent patients ^[7]^. Because surgical specimens reflect the cumulative neoadjuvant sequence (systemic chemotherapy, ablative SBRT, parenchymal debulking, and fibromesenchymal remodeling), transcriptional differences reflect treatment-associated tissue remodeling and cannot be attributed exclusively or cell-autonomously to radiation.

Differential expression statistics, pathological treatment-response classifications (responders vs. non-responders), and clinical annotations from the primary study were incorporated into PDAC BioPortal. Within the multimodal framework of the platform, this cohort provides a crucial clinical benchmark for evaluating whether candidate biomarkers and functional programs identified in untreated disease remain active, are extinguished, or undergo phenotypic reorganization following multimodal neoadjuvant therapy.

Single-nucleus RNA-sequencing (snRNA-seq) data were obtained from the human PDAC reference atlas deposited under GEO accession GSE202051 ^[5]^. The primary study generated 224,988 high-quality single-nucleus transcriptomic profiles from 43 surgical PDAC specimens, establishing a high-resolution molecular taxonomy of malignant epithelial cells, normal ductal epithelium, cancer-associated fibroblast (CAF) lineages, vascular endothelium, pericytes, and diverse immune compartments. The full cohort comprises 18 treatment-naïve resections and 25 resections following neoadjuvant therapy. All specimens were acquired at definitive surgical resection and represent independent patient donors without longitudinal pre/post pairing.

Within the neoadjuvant treated group (n = 25), the primary clinical backbone consisted of multi-cycle induction FOLFIRINOX followed by fractionated radiotherapy with concurrent fluoropyrimidine (5-fluorouracil or capecitabine; defined as “Standard CRT”, n = 14), with additional patient subsets receiving chemoradiotherapy combined with losartan or trial-based immunotherapy. To ensure therapeutic homogeneity and eliminate pharmacological confounding in comparative pathway analyses, cross-modal evaluations with the bulk SBRT cohort were restricted to the well-defined Standard CRT subset (n = 14) versus treatment-naïve controls (n = 18; 32 independent biological replicates), while the full 43-patient cohort was preserved for interactive single-nucleus visualization.

Cell-type annotations, cell metadata, and uniform manifold approximation and projection (UMAP) coordinates defined by the original study were retained without computational re-clustering. To bridge tissue-level bulk measurements with cellular resolution, PDAC BioPortal integrates two complementary analytical layers consisting of UMAP embedding, pseudobulk deconvolution, and GSEA pathway analysis.

#### 4.2.3. Spatial transcriptomic PDAC datasets

Spatial transcriptomic data were obtained from GEO accession GSE274103 ^[21]^, comprising five treatment-naïve formalin-fixed paraffin-embedded (FFPE) PDAC sections profiled on the 10x Genomics Visium Spatial Gene Expression platform ^[37]^. Following quality filtering, the integrated spatial dataset encompasses 23,436 tissue spots (55 μm spot diameter) and 17,943 measured genes across five independent clinical sections.

Raw spatial transcript counts were normalized using a counts-per-10,000 (CP10k) library-size scaling followed by a log⁡1p transformation. Normalized expression matrices, spot spatial coordinates, and corresponding high-resolution hematoxylin and eosin (H&E) brightfield histological images were converted into binary tiled assets for web deployment. During interactive querying, expression intensities for selected candidate genes are mapped directly onto their corresponding tissue coordinates and overlaid on the histological image. This module serves as an anatomical validation framework, enabling users to evaluate whether candidate genes identified in bulk or single-nucleus analyses exhibit focal localization within neoplastic ductal foci, desmoplastic stroma, vascular beds, or tertiary lymphoid structures.

### 4.3. Differential Expression Analysis and Statistical Methods

To minimize cross-platform batch artifacts while enabling consistent biological interpretation, differential expression analyses were executed independently within each cohort using statistical frameworks tailored to the original study designs. Direct pooling or uncalibrated cross-platform merging of absolute expression values between disparate sequencing platforms was avoided.

#### 4.3.1. TCGA-GTEx differential expression analysis

Tumor-versus-normal differential expression in the Toil-recomputed TCGA-GTEx matrix ^[35,36,38,39]^ compared 178 primary TCGA-PAAD tumors against 167 GTEx normal pancreas specimens (n = 345).

The primary differential expression analysis employed a two-sided Wilcoxon rank-sum test (Mann-Whitney U test) implemented via SciPy ^[40]^ to accommodate non-normal expression distributions in cross-cohort tissue samples. For each gene, the pipeline calculates the mean log2 fold change (log2FC), asymptotic p-value, Benjamini-Hochberg false discovery rate (FDR) ^[41]^, and rank-biserial correlation as a non-parametric effect size. By default, genes satisfying FDR < 0.05 and |log2FC| ≥ 1.0 were designated as significantly differentially expressed.

To provide parametric validation, a secondary workflow was performed on the corresponding RSEM expected-count matrix. Expected counts were rounded to integers and modeled using the Bioconductor edgeR ^[42]^ and limma-voom framework ^[43,44]^. Low-abundance transcripts were filtered using filterByExpr, library sizes were normalized via the Trimmed Mean of M-values (TMM) method, precision weights were estimated using voom, and moderated t-statistics were derived via empirical Bayes linear modeling (lmFit and eBayes). Genes exhibiting concordant significance across both Wilcoxon and limma-voom pipelines were prioritized as high-confidence disease-associated alterations.

#### 4.3.2. SBRT cohort analysis

The neoadjuvant radiotherapy module incorporates differential expression results from the GSE225767 study ^[7]^. Gene-level comparisons between pre-treatment biopsies (n = 26) and post-SBRT resections (n = 29) across 19,701 expressed genes were modeled using the negative binomial generalized linear model implemented in DESeq2 ^[45]^, with Wald test p-values adjusted using the Benjamini-Hochberg procedure ^[41]^. The portal maintains the published effect sizes, adjusted p-values, and pathological treatment-response classifications without re-estimation, ensuring exact fidelity to the original clinical trial dataset while enabling interactive exploration. For targeted interrogation of the curated 20-gene NRF2 panel on normalized log2 expression values, two-sided Mann-Whitney U tests were performed alongside panel-wide Benjamini-Hochberg FDR q-values across the 20 tested genes.

#### 4.3.3. Gene and pathway-level analyses

To advance interpretation from individual candidate genes to coordinated biological programs, pathway enrichment analysis was implemented across each transcriptomic cohort using the Molecular Signatures Database (MSigDB) Hallmark (50 gene sets) ^[46]^, Reactome (1,787 pathways), and Gene Ontology Biological Process (GO BP; 5,393 terms) collections, together with complementary over-representation analysis (ORA) of significantly differentially expressed genes via right-tailed hypergeometric testing with Benjamini-Hochberg FDR correction. Genes were ranked across the full analyzed background universe (19,853 genes in TCGA-GTEx; 19,701 genes in GSE225767) by the signed significance metric: Rank = sign(log2FC) × −log10(p-value). In interactive portal pathway analyses (Figures 2–3 and Supplementary Figure S1B-C), Kolmogorov-Smirnov running enrichment scores and leading-edge gene sets were computed directly across the ranked list, and enrichment magnitude was quantified via an analytical Mann-Whitney U rank-sum enrichment Z-score (ZGSEA; displayed as NES in the portal UI) with normal-tail p-values bounded at a numerical floor of p ≥ 1.00 × 10-6 (corresponding to Benjamini-Hochberg floors of FDR < 1.71 × 10-5 in TCGA-GTEx and FDR < 1.31 × 10-4 in GSE225767). For cross-cohort and cross-modal Hallmark pathway concordance analyses (Sections 4.5 and 2.2), classical pre-ranked GSEA ^[47]^ was executed using 1,000 gene-set permutations to yield permutation-normalized NES values (NESperm). Shared leading-edge core genes were extracted to evaluate pathway network overlap and biological cross-talk using Jaccard similarity indices. For multi-gene signature analyses, sample-level composite scores were computed as the mean of gene-wise Z-scores across each cohort and compared using Pearson (r) and Spearman (ρ) correlation analyses. Overall survival analyses in TCGA-PAAD (N = 177 primary tumors with complete clinical follow-up) were evaluated both as a continuous predictor via univariate Cox proportional hazards regression (reporting HR per unit signature Z-score, two-sided Wald p-value, and Schoenfeld residual proportional-hazards testing) and via exploratory optimal-cutpoint Kaplan-Meier stratification. For binary stratification, optimal cutpoints were selected via maximally selected Mantel-Cox log-rank statistics across the 15th-85th percentile range (retaining at least 15% of patients in each group), Kaplan-Meier curves with Greenwood 95% confidence intervals were estimated, and both the unadjusted log-rank p-value and the Miller-Siegmund maximally selected rank statistic adjusted p-value (maxstat padj) were calculated alongside Peto/Cox hazard ratios (HR) comparing High versus Low signature expression (where HR > 1.0 indicates increased hazard and shorter overall survival in the High-expression group).

### 4.4. Single-nucleus pseudobulk analysis

The single-nucleus atlas (GSE202051) ^[5]^ comprises 224,988 nuclei from 43 resected PDAC specimens: 18 treatment-naïve (108,964 nuclei) and 25 neoadjuvant-treated (116,024 nuclei), including 14 receiving standard fluoropyrimidine chemoradiotherapy (CRT) after FOLFIRINOX alongside CRT combinations with losartan, nivolumab, or investigational agents.

For statistical analyses and cross-modal comparison with bulk RNA-seq datasets, nuclei were aggregated into patient-level pseudobulk profiles to preserve the patient as the biological unit of replication and avoid pseudo-replication arising from treating individual nuclei as independent observations. Pseudobulk expression profiles were generated for whole tumor specimens and separately for major cellular compartments, including malignant epithelial cells, normal ductal cells, cancer-associated fibroblasts (CAFs), myofibroblastic CAFs (myCAFs), vascular endothelial cells, macrophages, and CD8+ T cells. This approach provided up to 43 biological replicates across the complete cohort and 32 biological replicates for the prespecified comparison of treatment-naïve tumors (n = 18) with tumors from patients receiving standard CRT (n = 14). For interactive UMAP visualization and efficient cross-cohort exploration within PDAC BioPortal, we additionally generated an evenly stratified subset of 20,000 nuclei comprising 9,689 treatment-naïve and 10,311 neoadjuvant-treated nuclei, while preserving the distribution of patients and major cellular lineages represented in the full dataset. Importantly, this visualization subset was used for interactive visualization rather than as a replacement for the full cohort in patient-level statistical analyses.

Two complementary statistical workflows were applied to the patient-level pseudobulk profiles. First, for gene-level sub-lineage comparisons (Section 2.3.3 and Supplementary Figure S2), patient-level pseudobulk mean expression (log1p CP10K) was computed per patient within each annotated sub-lineage and compared between treatment-naïve patients (n = 18) and either the full neoadjuvant-treated cohort (n = 25; up to 43 patients) or the standard CRT subgroup (n = 14; up to 32 patients) using two-sided Welch’s t-tests and Mann-Whitney U tests, with Benjamini-Hochberg FDR correction applied across evaluated sub-lineages. Second, for whole-sample and compartment-resolved pathway enrichment (Sections 4.5 and 2.2), aggregated patient-level pseudobulk counts comparing standard CRT (n = 14) versus treatment-naïve (n = 18) tumors were modeled on log2(CPM + 0.5) values using the empirical Bayes limma-trend framework ^[44]^, and the resulting moderated t-statistics across 22,164 genes were queried against the MSigDB Hallmark collection via 1,000-permutation pre-ranked GSEA ^[46,47]^.

### 4.5. Cross-modal treatment-associated pathway concordance and cellular source analysis

To test the reproducibility and generalizability of treatment-associated transcriptional programs between independent neoadjuvant cohorts, curated expression matrices and pre-computed Hallmark pathway enrichment profiles were downloaded directly from PDAC BioPortal. The portal’s standardized analytical exports provided harmonized differential expression outputs, normalized pseudobulk matrices, and cell-type metadata for both the bulk neoadjuvant SBRT cohort (GSE225767; ^[7]^) and the single-nucleus chemoradiotherapy reference atlas (GSE202051;^[5]^).

Using the processed data retrieved from PDAC BioPortal, cross-modal concordance was evaluated in three sequential steps:

#### 4.5.1. Global Pathway Rank Concordance and Compartment Deconvolution

Concordance across the 50 MSigDB Hallmark pathways was evaluated between bulk SBRT (GSE225767) and whole-sample single-nucleus CRT pseudobulk (GSE202051) using Spearman (ρ), Pearson (r), and directional quadrant mapping. To resolve cellular provenance, bulk NESperm values were cross-referenced against compartment-resolved NESperm profiles across malignant ductal cells, CAFs, myCAFs, vascular endothelium, macrophages, and CD8+ T cells.

#### 4.5.2. Cellular Composition Shift Analysis and Kitagawa-Oaxaca-Blinder Decomposition

Treatment-associated shifts in patient-level cell-type fractions between treatment-naïve (n = 18) and standard CRT (n = 14) specimens were evaluated across the nine major annotated lineages using two-sided Welch’s t-tests and Mann-Whitney U tests with Benjamini-Hochberg FDR correction. To quantitatively separate compositional remodeling from cell-intrinsic transcriptional adaptation, we applied a two-fold Kitagawa-Oaxaca-Blinder decomposition partitioning each whole-sample pseudobulk pathway shift into a within-cell-type expression component and a cellular-composition component: ΔȲg = Σk p̄k Δȳk,g + Σk ȳk,g Δpk, where p̄k and ȳk,g denote the cross-group mean fraction and mean expression of lineage k, and Δpk and Δȳk,g denote the treatment-minus-naïve changes in lineage proportion and intrinsic lineage expression, respectively. Because single-nucleus cell-type proportions lie on a closed simplex where depletion of one compartment mechanically inflates remaining fractions, we additionally performed a centered log-ratio (CLR) compositional sensitivity analysis across all annotated lineages (0.5-count pseudocount followed by two-sided Welch’s t-tests and BH FDR correction on CLR-transformed proportions).

### 4.6. PDACopilot and hierarchical model routing

PDACopilot was developed as an evidence-grounded natural-language interface to streamline exploration of the harmonized datasets within PDAC BioPortal. To ensure scientific fidelity, queries are classified by analytical intent, computational complexity, and evidence requirements, and processed through a three-tiered hierarchical routing architecture ^[22,23]^:

Level 0 / Tier 1 (Deterministic BioPortal Grounded Engine): Queries seeking exact numerical values, differential expression tables, GSEA statistics, or dataset metadata bypass generative language models entirely and are resolved directly via deterministic database queries, mitigating numerical hallucination by grounding quantitative outputs in deterministic queries.

Level 1 / Tier 2 (Conversational Navigation): Standard biological inquiries and navigational tasks are routed to Llama 3.3 70B via high-throughput inference engines.

Level 2 / Tier 3 (Complex Synthesis & Hypothesis Generation): Multi-step cross-cohort queries and cross-modal synthesis are directed to Google Gemini Flash.

In all language-model-assisted tiers, prompts are bound by strict retrieval-augmented contracts that force models to cite specific dataset IDs, sample sizes, and statistical confidence intervals retrieved by the Grounded Engine. The copilot is deployed via serverless Next.js API routes and edge worker proxies, keeping API credentials secure outside the client browser. To evaluate evidence fidelity and resistance to hallucination, PDACopilot was assessed on a frozen 20-question, 40-point scientific diagnostic benchmark (scored 0-2 points per query across four criteria: dataset routing correctness, empirical numerical accuracy, FDR interpretation, and study-design/causality adherence) covering tumor-versus-normal differential expression, GSE225767 SBRT comparisons, single-nucleus lineage localization, spatial transcriptomics, GSEA/ORA pathway analyses, and causal-overclaim guardrails, audited via deterministic trace verification (EvidenceValidator.ts).

### 4.7. Web implementation

PDAC BioPortal was built with Next.js (v16), React (v19), TypeScript, Tailwind CSS, Recharts, and HTML5 Canvas as a static client-side web application requiring no dedicated analytical backend. Version 1.2.0 corresponds to the frozen manuscript software release archived on Zenodo, with subsequent survival and signature extensions available in the live portal.

## Declarations

## Ethics statement

This study analyzed exclusively publicly available, de-identified human transcriptomic datasets obtained from The Cancer Genome Atlas (TCGA), the Genotype-Tissue Expression (GTEx) project, and the NCBI Gene Expression Omnibus (GEO: GSE225767, GSE202051, and GSE274103). No new human subjects were recruited, and no additional institutional review board (IRB) approval or informed consent was required.

## Data availability

The transcriptomic datasets incorporated into PDAC BioPortal were obtained from publicly available resources. Harmonized bulk RNA-sequencing data for primary TCGA-PAAD tumors, TCGA adjacent normal specimens, and GTEx normal pancreas are available through the UCSC Xena Toil Recompute repository (TCGA-TARGET-GTEx cohort; https://xenabrowser.net/). The neoadjuvant treatment-associated bulk RNA-sequencing dataset is available through the Gene Expression Omnibus (GEO) under accession GSE225767. The single-nucleus PDAC reference atlas and spatial transcriptomic dataset are available through GEO under accessions GSE202051 and GSE274103, respectively. The corresponding source information and dataset provenance are retained within the portal. The processed portal release, associated metadata, and software archive are publicly available through Zenodo (DOI: 10.5281/zenodo.22149338). The PDAC BioPortal web application is publicly accessible through the project website: https://limboper89.github.io/PAAD-SBRT-GEx-Dashboard/.

## Code availability

The source code for PDAC BioPortal and PDACopilot, corresponding to release version 1.2.0 archived on Zenodo (DOI: 10.5281/zenodo.22149338), is publicly available through the PAAD-SBRT-GEx-Dashboard GitHub repository (https://github.com/Limboper89/PAAD-SBRT-GEx-Dashboard).

## CRediT author statement

Prince Kumar: Conceptualization, Methodology, Software, Data Curation, Formal Analysis, Visualization, Writing - Original Draft, Writing - Review & Editing. Erina Vlashi: Conceptualization, Supervision, Resources, Funding Acquisition, Writing - Review & Editing. Both authors read and approved the final manuscript.

## Competing interests

The authors declare no competing financial or non-financial interests.

## Funding

This work was supported by departmental research funds from the Department of Radiation Oncology, University of California, Los Angeles (UCLA).

## Acknowledgments

The authors thank the investigators, patients, and consortia who generated and made publicly accessible the TCGA-PAAD, GTEx, UCSC Xena Toil Recompute, GSE225767, GSE202051, and GSE274103 datasets.

## Declaration of generative AI use

In addition to the PDACopilot software module evaluated as an integral component of this study, generative AI language tools were used solely to assist with manuscript language polishing and formatting; all scientific analyses, statistical outputs, biological interpretations, and final text were verified and approved by the authors.

## Source Code & Archive

https://github.com/Limboper89/PAAD-SBRT-GEx-Dashboard | Zenodo DOI: 10.5281/zenodo.22149338

---

**Supplementary Figure S1.** Pathway-level NRF2 enrichment, ferroptosis survival analysis, and Radiation Sensitivity Index (RSI) correlations. (A) Exploratory Kaplan-Meier plot for the three-gene Iron/Ferroptosis signature in TCGA-PAAD (N = 177; continuous Cox p = 0.630; Table S6). (B) Rank-sum enrichment plot for Reactome KEAP1-NFE2L2 signaling in TCGA-GTEx (ZGSEA = +8.40, FDR < 1.71 × 10⁻⁵). (C) Enrichment plot for GO cellular response to oxidative stress in GSE225767 (ZGSEA = +2.07, FDR = 0.131). (D-E) Correlations between the eight-gene NRF2 target signature and 10-gene RSI in TCGA-PAAD (n = 178; r = 0.869, p = 1.15 × 10⁻⁵⁵; portal UI clamped at 1.00 × 10⁻⁹⁹) and GSE225767 (N = 55; r = 0.743, p = 8.28 × 10⁻¹¹).

**Supplementary Figure S2.** Patient-level pseudobulk comparison of NFE2L2 expression across PDAC cellular sub-lineages. Mean patient-level pseudobulk NFE2L2 expression (log1p ± SE) comparing treatment-naïve patients (n = 18, teal) against (A) the full neoadjuvant-treated cohort (n = 25, orange; FDR q = 0.027 for dendritic, delta, and vascular endothelial cells; q = 0.047 for alpha cells; nominal p = 0.02 for malignant epithelial cells) and (B) the standard chemoradiotherapy (CRT) subgroup (n = 14, orange). Patient sample counts per compartment (N/T) are indicated below each sub-lineage.

**Supplementary Figure S3.** Cost-aware, evidence-grounded hierarchical routing architecture of PDACopilot. (A-F) Execution pipeline from natural-language user query (A) and ModelRouter intent/complexity classification (B) through three hierarchical resolution tiers (C-D): Level 0 (Deterministic BioPortal Grounded Engine), Level 1 (Llama 3.3 70B lightweight interpretation), and Level 2 (Google Gemini Flash complex cross-modal synthesis), followed by EvidenceValidator consistency gating (E) and delivery of traceable, dataset-cited responses (F). Lower panels summarize deterministic evidence grounding, design principles, deployment topologies (Next.js server API route and Cloudflare Worker proxy), and API key security.

## Notes

https://limboper89.github.io/PAAD-SBRT-GEx-Dashboard/

https://doi.org/10.5281/zenodo.22149338

### Competing Interest Statement

The authors have declared no competing interest.

### Summary of Updates

The previous abstract was 400 words. The abstract has been cut down to 176 words. The manuscript was extremely wordy which has been cut down by separating some data to supplementary tables. Also manuscript file was in non standard formatting which has been substantially improved and included in current version.

## References

1. Hosein AN, Brekken RA, Maitra A. Pancreatic cancer stroma: an update on therapeutic targeting strategies. Nat Rev Gastroenterol Hepatol. 2020;17(8):487–505. doi:10.1038/s41575-020-0300-1. PMID:32393771.

2. Siegel RL, Giaquinto AN, Jemal A. Cancer statistics, 2024. CA Cancer J Clin. 2024;74(1):12–49. doi:10.3322/caac.21820. PMID:38230766.

3. Bailey P, Chang DK, Nones K, et al. Genomic analyses identify molecular subtypes of pancreatic cancer. Nature. 2016;531(7592):47–52. doi:10.1038/nature16965. PMID:26909576.

4. Biffi G, Oni TE, Spielman B, et al. IL1-Induced JAK/STAT Signaling Is Antagonized by TGFβ to Shape CAF Heterogeneity in Pancreatic Ductal Adenocarcinoma. Cancer Discov. 2019;9(2):282–301. doi:10.1158/2159-8290.CD-18-0710. PMID:30366930.

5. Hwang WL, Jagadeesh KA, Guo JA, et al. Single-nucleus and spatial transcriptome profiling of pancreatic cancer identifies multicellular dynamics associated with neoadjuvant treatment. Nat Genet. 2022;54(8):1178–1191. doi:10.1038/s41588-022-01134-8. PMID:35902743.

6. Grünwald BT, Devisme A, Andrieux G, et al. Spatially confined sub-tumor microenvironments in pancreatic cancer. Cell. 2021;184(22):5577–5592.e18. doi:10.1016/j.cell.2021.09.022. PMID:34644529.

7. Piper M, Hoen M, Darragh LB, et al. Simultaneous targeting of PD-1 and IL-2Rβγ with radiation therapy inhibits pancreatic cancer growth and metastasis. Cancer Cell. 2023;41(5):950–969.e6. doi:10.1016/j.ccell.2023.04.001. PMID:37116489.

8. Benitz S, Steep A, Nasser MM, et al. ROR2 Regulates Cellular Plasticity in Pancreatic Neoplasia and Adenocarcinoma. Cancer Discov. 2024;14(11):2162–2182. doi:10.1158/2159-8290.CD-24-0137. PMID:38975886.

9. Cancer Genome Atlas Research Network. Integrated Genomic Characterization of Pancreatic Ductal Adenocarcinoma. Cancer Cell. 2017;32(2):185–203.e13. doi:10.1016/j.ccell.2017.07.007. PMID:28810144.

10. Shiau C, Su J, Guo JA, et al. Treatment-associated remodeling of the pancreatic cancer endothelium at single-cell resolution. Front Oncol. 2022;12:929950. doi:10.3389/fonc.2022.929950. PMID:36185212.

11. Tang Z, Kang B, Li C, et al. GEPIA2: an enhanced web server for large-scale expression profiling and interactive analysis. Nucleic Acids Res. 2019;47(W1):W556–W560. doi:10.1093/nar/gkz430. PMID:31114875.

12. Cerami E, Gao J, Dogrusoz U, et al. The cBio cancer genomics portal: an open platform for exploring multidimensional cancer genomics data. Cancer Discov. 2012;2(5):401–404. doi:10.1158/2159-8290.CD-12-0095. PMID:22588877.

13. Gao J, Aksoy BA, Dogrusoz U, et al. Integrative analysis of complex cancer genomics and clinical profiles using the cBioPortal. Sci Signal. 2013;6(269):pl1. doi:10.1126/scisignal.2004088. PMID:23550210.

14. Han Y, Wang Y, Dong X, et al. TISCH2: expanded datasets and new tools for single-cell transcriptome analyses of the tumor microenvironment. Nucleic Acids Res. 2023;51(D1):D1425–D1431. doi:10.1093/nar/gkac959. PMID:36321662.

15. Megill C, Martin B, Weaver C, et al. cellxgene: a performant, scalable exploration platform for high dimensional sparse matrices. bioRxiv. 2021:2021.04.05.438318. doi:10.1101/2021.04.05.438318.

16. Chen Z, Luo Z, Zhang D, et al. TIGER: A Web Portal of Tumor Immunotherapy Gene Expression Resource. Genomics Proteomics Bioinformatics. 2023;21(2):337–348. doi:10.1016/j.gpb.2022.08.004. PMID:36049666.

17. Maurer C, Holmstrom SR, He J, et al. Experimental microdissection enables functional harmonisation of pancreatic cancer subtypes. Gut. 2019;68(6):1034–1043. doi:10.1136/gutjnl-2018-317706. PMID:30658994.

18. Versteijne E, Suker M, Groothuis K, et al. Preoperative Chemoradiotherapy Versus Immediate Surgery for Resectable and Borderline Resectable Pancreatic Cancer: Results of the Dutch Randomized Phase III PREOPANC Trial. J Clin Oncol. 2020;38(16):1763–1773. doi:10.1200/JCO.19.02274. PMID:32105518.

19. Katz MHG, Shi Q, Meyers J, et al. Efficacy of Preoperative mFOLFIRINOX vs mFOLFIRINOX Plus Hypofractionated Radiotherapy for Borderline Resectable Adenocarcinoma of the Pancreas: The A021501 Phase 2 Randomized Clinical Trial. JAMA Oncol. 2022;8(9):1263–1270. doi:10.1001/jamaoncol.2022.2319. PMID:35834226.

20. GTEx Consortium. The GTEx Consortium atlas of genetic regulatory effects across human tissues. Science. 2020;369(6509):1318–1330. doi:10.1126/science.aaz1776. PMID:32913098.

21. Liu Y, Sinjab A, Min J, et al. Conserved spatial subtypes and cellular neighborhoods of cancer-associated fibroblasts revealed by single-cell spatial multi-omics. Cancer Cell. 2025;43(5):905–924.e6. doi:10.1016/j.ccell.2025.03.004. PMID:40154487.

22. Moor M, Banerjee O, Abad ZSH, et al. Foundation models for generalist medical artificial intelligence. Nature. 2023;616(7956):259–265. doi:10.1038/s41586-023-05881-4. PMID:37045921.

23. Jin Q, Yang Y, Chen Q, et al. GeneGPT: augmenting large language models with domain tools for improved access to biomedical information. Bioinformatics. 2024;40(2):btae075. doi:10.1093/bioinformatics/btae075. PMID:38341654.

24. DeNicola GM, Karreth FA, Humpton TJ, et al. Oncogene-induced Nrf2 transcription promotes ROS detoxification and tumorigenesis. Nature. 2011;475(7354):106–109. doi:10.1038/nature10189. PMID:21734707.

25. Chio IIC, Jafarnejad SM, Ponz-Sarvise M, et al. NRF2 Promotes Tumor Maintenance by Modulating mRNA Translation in Pancreatic Cancer. Cell. 2016;166(4):963–976. doi:10.1016/j.cell.2016.06.056. PMID:27477511.

26. Rojo de la Vega M, Chapman E, Zhang DD. NRF2 and the Hallmarks of Cancer. Cancer Cell. 2018;34(1):21–43. doi:10.1016/j.ccell.2018.03.022. PMID:29731393.

27. Grass GD, Alfonso JCL, Welsh E, et al. The Radiosensitivity Index Gene Signature Identifies Distinct Tumor Immune Microenvironment Characteristics Associated With Susceptibility to Radiation Therapy. Int J Radiat Oncol Biol Phys. 2022 Jul 1;113(3):635–647. doi: 10.1016/j.ijrobp.2022.03.006. Epub 2022 Mar 12. PMID: 35289298; PMCID: PMC9250625.

28. Dixon SJ, Lemberg KM, Lamprecht MR, et al. Ferroptosis: an iron-dependent form of nonapoptotic cell death. Cell. 2012;149(5):1060–1072. doi:10.1016/j.cell.2012.03.042.

29. Yang WS, SriRamaratnam R, Welsch ME, et al. Regulation of ferroptotic cancer cell death by GPX4. Cell. 2014;156(1-2):317–331. doi:10.1016/j.cell.2013.12.010.

30. Dixon SJ, Olzmann JA. The cell biology of ferroptosis. Nature Reviews Molecular Cell Biology. 2024;25(6):424–442. doi:10.1038/s41580-024-00703-5.

31. Dixon SJ, Stockwell BR. The Hallmarks of Ferroptosis. Annu Rev Cancer Biol. 2019;3:35–54. doi:10.1146/annurev-cancerbio-030518-055844.

32. Clusmann J, Kolbinger FR, Muti HS, et al. The future landscape of large language models in medicine. Commun Med (Lond). 2023;3(1):141. doi:10.1038/s43856-023-00370-1. PMID:37816837.

33. Thirunavukarasu AJ, Ting DSJ, Elangovan K, et al. Large language models in medicine. Nat Med. 2023;29(8):1930–1940. doi:10.1038/s41591-023-02448-8. PMID:37460753.

34. Lewis P, Perez E, Piktus A, et al. Retrieval-Augmented Generation for Knowledge-Intensive NLP Tasks. In: Advances in Neural Information Processing Systems (NeurIPS 2020). 2020;33:9459–9474. doi:10.48550/arXiv.2005.11401.

35. Goldman MJ, Craft B, Hastie M, et al. Visualizing and interpreting cancer genomics data via the Xena platform. Nat Biotechnol. 2020;38(6):675–678. doi:10.1038/s41587-020-0546-8. PMID:32444850.

36. Vivian J, Rao AA, Nothaft FA, et al. Toil enables reproducible, open source, big biomedical data analyses. Nat Biotechnol. 2017;35(4):314–316. doi:10.1038/nbt.3772. PMID:28398314.

37. Ståhl PL, Salmén F, Vickovic S, et al. Visualization and analysis of gene expression in tissue sections by spatial transcriptomics. Science. 2016;353(6294):78–82. doi:10.1126/science.aaf2403. PMID:27365449.

38. Dobin A, Davis CA, Schlesinger F, et al. STAR: ultrafast universal RNA-seq aligner. Bioinformatics. 2013;29(1):15–21. doi:10.1093/bioinformatics/bts635. PMID:23104886.

39. Li B, Dewey CN. RSEM: accurate transcript quantification from RNA-Seq data with or without a reference genome. BMC Bioinformatics. 2011;12:323. doi:10.1186/1471-2105-12-323. PMID:21816040.

40. Virtanen P, Gommers R, Oliphant TE, et al. SciPy 1.0: fundamental algorithms for scientific computing in Python. Nat Methods. 2020;17(3):261–272. doi:10.1038/s41592-019-0686-2. PMID:32015543.

41. Benjamini Y, Hochberg Y. Controlling the False Discovery Rate: A Practical and Powerful Approach to Multiple Testing. J R Stat Soc Series B. 1995;57(1):289–300. doi:10.1111/j.2517-6161.1995.tb02031.x.

42. Robinson MD, McCarthy DJ, Smyth GK. edgeR: a Bioconductor package for differential expression analysis of digital gene expression data. Bioinformatics. 2010;26(1):139–140. doi:10.1093/bioinformatics/btp616. PMID:19910308.

43. Law CW, Chen Y, Shi W, et al. voom: Precision weights unlock linear model analysis tools for RNA-seq read counts. Genome Biol. 2014;15(2):R29. doi:10.1186/gb-2014-15-2-r29. PMID:24485249.

44. Ritchie ME, Phipson B, Wu D, et al. limma powers differential expression analyses for RNA-sequencing and microarray studies. Nucleic Acids Res. 2015;43(7):e47. doi:10.1093/nar/gkv007. PMID:25605792.

45. Love MI, Huber W, Anders S. Moderated estimation of fold change and dispersion for RNA-seq data with DESeq2. Genome Biol. 2014;15(12):550. doi:10.1186/s13059-014-0550-8. PMID:25516281.

46. Liberzon A, Birger C, Thorvaldsdóttir H, et al. The Molecular Signatures Database (MSigDB) hallmark gene set collection. Cell Syst. 2015;1(6):417–425. doi:10.1016/j.cels.2015.12.004. PMID:26771021.

47. Subramanian A, Tamayo P, Mootha VK, et al. Gene set enrichment analysis: a knowledge-based approach for interpreting genome-wide expression profiles. Proc Natl Acad Sci U S A. 2005;102(43):15545–15550. doi:10.1073/pnas.0506580102. PMID:16199517.

