## Supplementary tables for "PDAC BioPortal: An Integrated Web Resource for Multi-Scale Analysis of Pancreatic Cancer Transcriptomics"

### Supplementary Tables: PDAC BioPortal

**Supplementary Table S1. Differential expression of the 20-gene NRF2-associated redox panel in primary PDAC (TCGA-PAAD vs. GTEx) and neoadjuvant treatment-associated PDAC (GSE225767).**

| Gene | Functional Module | TCGA-GTEx log2FC | TCGA-GTEx FDR q | GSE225767 Δ (log2) | MW p-value | 20-Gene BH q | DESeq2 log2FC | DESeq2 adj. p |
|---|---|---|---|---|---|---|---|---|
| NFE2L2 | Core NRF2 Machinery | +2.12 | 2.89e-50 | +0.73 | 0.0637 | 0.127 | +0.73 | 1.64e-01 |
| KEAP1 | Core NRF2 Machinery | +2.25 | 5.25e-54 | +1.42 | 0.3063 | 0.469 | +1.42 | 1.73e-01 |
| MAFF | Core NRF2 Machinery | +3.26 | 5.88e-51 | +0.95 | 0.0191 | 0.082 | +0.95 | 8.69e-02 |
| MAFG | Core NRF2 Machinery | +1.49 | 2.20e-43 | -0.16 | 0.7809 | 0.868 | -0.16 | 8.40e-01 |
| NQO1 | Canonical Antioxidant | +6.05 | 3.92e-54 | -0.64 | 0.0234 | 0.082 | -0.64 | 3.19e-01 |
| HMOX1 | Canonical Antioxidant | +3.64 | 3.13e-52 | +1.31 | 0.0303 | 0.082 | +1.31 | 1.82e-01 |
| PRDX1 | Canonical Antioxidant | +3.07 | 4.81e-54 | +0.41 | 0.3282 | 0.469 | +0.41 | 5.52e-01 |
| TXNRD1 | Canonical Antioxidant | +2.54 | 1.32e-51 | -0.38 | 0.6369 | 0.756 | -0.38 | 4.49e-01 |
| SLC7A11 | Glutathione & Cystine | +1.19 | 2.44e-18 | +1.52 | 0.0416 | 0.092 | +1.52 | 1.48e-01 |
| GCLC | Glutathione & Cystine | +0.80 | 8.94e-21 | +3.49 | 0.0017 | 0.035 | +3.49 | 6.64e-04 |
| GCLM | Glutathione & Cystine | +2.09 | 9.10e-47 | -0.14 | 0.6429 | 0.756 | -0.14 | 8.37e-01 |
| GSS | Glutathione & Cystine | +2.72 | 4.30e-54 | +1.02 | 0.0330 | 0.082 | +1.02 | 9.35e-02 |
| GSTP1 | Glutathione & Cystine | +2.48 | 1.79e-46 | -0.01 | 0.9664 | 0.966 | -0.01 | 1.00e+00 |
| ME1 | NADPH Metabolism | +1.87 | 1.19e-39 | +1.37 | 0.5379 | 0.717 | +1.37 | 2.40e-01 |
| IDH1 | NADPH Metabolism | +2.21 | 5.04e-51 | +1.69 | 0.2563 | 0.427 | +1.69 | 1.36e-01 |
| G6PD | NADPH Metabolism | +2.30 | 2.24e-52 | +0.56 | 0.2413 | 0.427 | +0.56 | 2.71e-01 |
| PGD | NADPH Metabolism | +2.63 | 1.05e-53 | -0.33 | 0.9127 | 0.961 | -0.33 | 5.49e-01 |

| Gene | Functional Module | TCGA-GTEx log2FC | TCGA-GTEx FDR q | GSE225767 Δ (log2) | MW p-value | 20-Gene BH q | DESeq2 log2FC | DESeq2 adj. p |
|---|---|---|---|---|---|---|---|---|
| FTH1 | Iron & Ferroptosis Defense | +1.32 | 5.17e-37 | +0.40 | 0.0290 | 0.082 | +0.40 | 1.32e-01 |
| FTL | Iron & Ferroptosis Defense | +2.74 | 5.85e-54 | +0.69 | 0.0106 | 0.082 | +0.69 | 4.45e-02 |
| GPX4 | Iron & Ferroptosis Defense | +2.83 | 7.11e-54 | +1.04 | 0.0132 | 0.082 | +1.04 | 5.73e-02 |

*Note: TCGA-GTEx evaluates n = 178 primary PDAC tumors vs. n = 167 GTEx normal pancreas via two-sided Wilcoxon rank-sum tests. GSE225767 compares unpaired pre-treatment biopsies (n = 26) vs. post-neoadjuvant surgical resections (n = 29); MW = Mann-Whitney U test; BH q = Benjamini-Hochberg FDR across the 20-gene panel.*

**Supplementary Table S2. Patient-level single-nucleus cellular composition shifts (GSE202051: n = 18 treatment-naïve vs. n = 14 standard CRT) with centered log-ratio (CLR) compositional sensitivity analysis.**

| Cell Lineage | Naïve Mean % | CRT Mean % | Δ % | Welch p | Welch FDR | MW p | MW FDR | CLR Welch p | CLR FDR |
|---|---|---|---|---|---|---|---|---|---|
| Malignant | 49.6% | 14.5% | -35.2% | 7.09e-04 | 0.0064 | 8.87e-04 | 0.0080 | 6.83e-04 | 0.0061 |
| CAF | 12.9% | 19.5% | +6.6% | 0.202 | 0.2622 | 0.279 | 0.4185 | 0.180 | 0.3445 |
| Ductal | 6.0% | 17.2% | +11.2% | 0.059 | 0.2622 | 0.143 | 0.3472 | 0.191 | 0.3445 |
| Vascular | 5.3% | 8.4% | +3.1% | 0.112 | 0.2622 | 0.154 | 0.3472 | 0.504 | 0.7566 |
| Pericyte | 2.8% | 7.0% | +4.2% | 0.182 | 0.2622 | 0.747 | 0.8643 | 0.685 | 0.8804 |
| myCAF | 5.9% | 2.0% | -3.8% | 0.132 | 0.2622 | 0.224 | 0.4038 | 0.160 | 0.3445 |
| Macrophage | 3.5% | 3.6% | +0.1% | 0.945 | 0.9450 | 0.864 | 0.8643 | 0.905 | 0.9048 |
| CD8+ T | 1.7% | 2.7% | +1.0% | 0.522 | 0.5874 | 0.833 | 0.8643 | 0.869 | 0.9048 |
| Acinar | 1.8% | 5.4% | +3.7% | 0.204 | 0.2622 | 0.077 | 0.3444 | 0.060 | 0.2709 |

*Note: CLR = centered log-ratio transformation across the closed cell-fraction simplex (0.5 count pseudocount). Malignant epithelial depletion remains significant across all parametric, non-parametric, and CLR compositional tests.*

**Supplementary Table S3. Spatial transcriptomic distribution of NFE2L2 across five treatment-naïve PDAC 10x Visium sections (GSE274103).**

| Patient ID | GEO Sample | Total Spots | NFE2L2+ Spots | Spatial Coverage (%) | Mean Intensity (log1p, +spots) | Max Intensity (log1p) |
|---|---|---|---|---|---|---|
| PDAC-p1 | GSM8443449 | 4,987 | 1,990 | 39.9% | 1.627 | 3.768 |
| PDAC-p2 | GSM8443450 | 4,380 | 2,494 | 56.9% | 1.142 | 3.146 |
| PDAC-p3 | GSM8443451 | 4,134 | 1,348 | 32.6% | 1.434 | 3.537 |
| PDAC-p4 | GSM8443452 | 4,983 | 2,520 | 50.6% | 1.356 | 3.807 |
| PDAC-p5 | GSM8443453 | 4,952 | 2,159 | 43.6% | 1.432 | 4.234 |
| Total / Overall | 5 Sections | 23,436 | 10,511 | 44.8% | 1.142-1.627 | 3.146-4.234 |

*Note: Raw spot counts were CP10k-normalized and log1p-transformed.*

**Supplementary Table S4. PDACopilot evidence-consistency benchmark across language models and routing configurations.**

| Model / Configuration | Questions | Score | Full (2/2) | Partial (1/2) | Fail (0/2) | Question Drift | Stat Errors | Causal Overclaims | Num. Hallucinations | Routing Errors |
|---|---|---|---|---|---|---|---|---|---|---|
| Llama 3.3 70B (v1.3) | 20 | 15.2 / 40 (38.1%) | 4 | 7 | 9 | 4 | 2 | 2 | 1 | 2 |
| Gemini 2.5 Flash (Baseline, Unconstrained) | 20 | 20.0 / 40 (50.0%) | 6 | 9 | 5 | 3 | 2 | 2 | 1 | 2 |
| Gemini 2.5 Flash (Evidence-Locked v1.3+) | 20 | 39.0 / 40 (97.5%) | 19 | 1 | 0 | 0 | 0 | 0 | 0 | 0 |
| Gemini 2.5 Flash (Expanded 10-Criterion Suite) | 35 | 342 / 350 (97.7%) | - | - | - | 0 | 0 | 0 | 0 | 0 |

*Note: Baseline Gemini 2.5 Flash failures included FDR significance inversions on PHGDH (Q6, Q18), unpaired-cohort overclaim on GSE225767 (Q11), and a fabricated EPCAM p-value table (Q13). Expanded 35-question suite evaluated 10 criteria (0-10 points per question).*

**Supplementary Table S5. Cross-modal MSigDB Hallmark pathway concordance (bulk GSE225767 vs. single-nucleus pseudobulk GSE202051), Kitagawa-Oaxaca-Blinder decomposition, and compartment-resolved NESperm values.**

| Hallmark Pathway | Bulk NESperm | Bulk FDR | snRNA NESperm | snRNA FDR | Comp. % | Within-Cell % | Malignant NES | CAF NES | Vascular NES | CD8+ T NES |
|---|---|---|---|---|---|---|---|---|---|---|

| Hallmark Pathway | Bulk NESperm | Bulk FDR | snRNA NESperm | snRNA FDR | Comp. % | Within-Cell % | Malignant NES | CAF NES | Vascular NES | CD8+ T NES |
|---|---|---|---|---|---|---|---|---|---|---|
| TNF-alpha Signaling via NF-kB | +1.16 | 0.097 | -1.51 | 0.005 | 28.8% | 71.2% | -0.80 | +0.33 | +1.80 | +1.23 |
| Hypoxia | +1.24 | 0.009 | -1.85 | 0.002 | 5.3% | 94.7% | -0.78 | -0.96 | -1.02 | +1.10 |
| Cholesterol Homeostasis | +0.89 | 0.947 | -2.01 | 0.002 | 3.6% | 96.4% | -0.92 | -0.97 | -1.14 | +1.35 |
| Mitotic Spindle | +0.88 | 1.000 | -2.47 | 0.002 | 8.5% | 91.5% | -1.08 | -1.39 | -1.32 | +1.38 |
| Wnt-beta Catenin Signaling | +1.20 | 0.232 | -1.65 | 0.009 | 15.7% | 84.3% | -1.27 | -1.64 | -1.82 | +1.35 |
| TGF-beta Signaling | +1.24 | 0.134 | -2.12 | 0.002 | 10.4% | 89.6% | -1.22 | -1.29 | -1.65 | +1.10 |
| IL-6/JAK/STAT3 Signaling | +1.12 | 0.346 | -0.78 | 0.663 | 31.8% | 68.2% | -0.72 | +0.94 | +1.07 | -1.14 |
| DNA Repair | +0.91 | 0.947 | -2.07 | 0.002 | 38.2% | 61.8% | -0.96 | -1.04 | -0.98 | +0.99 |
| G2-M Checkpoint | +0.73 | 1.000 | -2.58 | 0.002 | 44.0% | 56.0% | -1.17 | -0.70 | -1.14 | +1.22 |
| Apoptosis | +1.25 | 0.009 | -1.59 | 0.002 | 20.9% | 79.1% | -1.00 | -0.85 | +1.11 | +1.05 |
| Notch Signaling | +1.32 | 0.103 | -1.87 | 0.002 | 38.1% | 61.9% | -1.19 | -1.25 | -1.04 | +1.43 |
| Adipogenesis | +1.17 | 0.077 | -1.84 | 0.002 | 14.5% | 85.5% | -1.13 | -1.13 | -0.94 | +1.09 |
| Estrogen Response Early | +1.15 | 0.128 | -1.91 | 0.002 | 11.5% | 88.5% | -0.73 | +0.26 | -0.90 | +1.07 |
| Estrogen Response Late | +1.08 | 0.346 | -2.14 | 0.002 | 29.4% | 70.6% | -0.72 | +1.08 | -1.07 | +1.08 |
| Androgen Response | +1.15 | 0.231 | -1.67 | 0.003 | 1.5% | 98.5% | -0.81 | -0.89 | -0.80 | +1.20 |
| Myogenesis | +1.52 | 0.006 | -1.07 | 0.314 | 22.8% | 77.2% | -0.98 | -0.98 | -1.01 | +1.12 |
| Protein Secretion | +0.93 | 0.859 | -2.00 | 0.002 | 8.1% | 91.9% | -1.04 | -1.55 | -0.89 | +0.97 |
| Interferon Alpha Response | +0.56 | 1.000 | -2.04 | 0.002 | 2.0% | 98.0% | -0.98 | -1.22 | +0.89 | -0.91 |
| Interferon Gamma Response | +1.14 | 0.152 | -1.68 | 0.002 | 9.0% | 91.0% | -0.96 | -0.86 | +1.18 | -0.88 |
| Apical Junction | +1.45 | 0.006 | -2.15 | 0.002 | 10.7% | 89.3% | -1.11 | -1.22 | -1.18 | +1.18 |
| Apical Surface | +1.07 | 0.522 | -1.39 | 0.066 | 11.5% | 88.5% | -1.12 | -1.16 | -1.00 | -1.10 |
| Hedgehog Signaling | +1.41 | 0.009 | -1.27 | 0.107 | 11.5% | 88.5% | -1.14 | -1.35 | -1.24 | +1.28 |
| Complement | +1.04 | 0.522 | -1.55 | 0.002 | 6.9% | 93.1% | -0.89 | -0.90 | -1.16 | +0.83 |

| Hallmark Pathway | Bulk NESperm | Bulk FDR | snRNA NESperm | snRNA FDR | Comp. % | Within-Cell % | Malignant NES | CAF NES | Vascular NES | CD8+ T NES |
|---|---|---|---|---|---|---|---|---|---|---|
| Unfolded Protein Response | +1.04 | 0.522 | -1.81 | 0.002 | 9.4% | 90.6% | -0.92 | -0.94 | -0.97 | +0.99 |
| PI3K/AKT/mTOR Signaling | +0.98 | 0.764 | -1.45 | 0.027 | 3.8% | 96.2% | -0.76 | -1.09 | -1.16 | +0.81 |
| mTORC1 Signaling | +0.84 | 1.000 | -2.16 | 0.002 | 30.7% | 69.3% | -0.76 | +0.21 | +1.09 | +1.30 |
| E2F Targets | +0.78 | 1.000 | -2.53 | 0.002 | 47.8% | 52.2% | -1.22 | -0.67 | -0.90 | +0.96 |
| Myc Targets V1 | +0.89 | 1.000 | -2.11 | 0.002 | 43.2% | 56.8% | -0.64 | +1.18 | -0.65 | +1.29 |
| Myc Targets V2 | +1.06 | 0.522 | -1.66 | 0.006 | 96.4% | 3.6% | -0.63 | +1.07 | -0.53 | +0.93 |
| Epithelial Mesenchymal Transition | +1.94 | 0.006 | -1.64 | 0.002 | 6.5% | 93.5% | -0.79 | -1.33 | -1.24 | +1.38 |
| Inflammatory Response | +1.15 | 0.128 | -0.97 | 0.461 | 35.8% | 64.2% | -0.78 | +0.23 | +1.45 | +0.87 |
| Xenobiotic Metabolism | +1.17 | 0.093 | -1.43 | 0.016 | 7.2% | 92.8% | -0.85 | -0.64 | -0.79 | +1.13 |
| Fatty Acid Metabolism | +1.02 | 0.594 | -1.60 | 0.002 | 6.6% | 93.4% | -0.94 | -1.08 | -1.11 | -0.92 |
| Oxidative Phosphorylation | +0.98 | 0.764 | -2.14 | 0.002 | 38.8% | 61.2% | -1.08 | -1.16 | -0.85 | +0.94 |
| Glycolysis | +0.97 | 0.806 | -2.37 | 0.002 | 3.2% | 96.8% | -0.71 | -1.06 | -0.96 | +1.41 |
| Reactive Oxygen Species Pathway | +0.73 | 1.000 | -1.82 | 0.003 | 0.9% | 99.1% | -0.62 | -0.73 | -0.74 | +0.92 |
| p53 Pathway | +1.12 | 0.248 | -1.94 | 0.002 | 21.0% | 79.0% | -0.80 | -0.82 | +1.26 | +0.90 |
| UV Response Up | +1.11 | 0.248 | -1.88 | 0.002 | 6.8% | 93.2% | -0.83 | -1.06 | -0.82 | +0.83 |
| UV Response Dn | +1.51 | 0.006 | -1.29 | 0.081 | 7.3% | 92.7% | -1.10 | -1.48 | -1.13 | +1.23 |
| Angiogenesis | +1.65 | 0.006 | -0.98 | 0.344 | 6.2% | 93.8% | -0.99 | -1.56 | -1.03 | +1.58 |
| heme Metabolism | -0.26 | 0.006 | -1.65 | 0.002 | 10.6% | 89.4% | -1.00 | -1.20 | -1.19 | +1.18 |
| Coagulation | +1.37 | 0.006 | -1.16 | 0.188 | 3.2% | 96.8% | -0.73 | -0.71 | -0.89 | +1.00 |
| IL-2/STAT5 Signaling | +1.23 | 0.013 | -1.26 | 0.084 | 13.2% | 86.8% | -0.78 | -0.92 | +1.07 | -0.92 |
| Bile Acid Metabolism | +1.07 | 0.483 | -1.10 | 0.258 | 15.2% | 84.8% | -1.15 | -1.04 | -0.82 | -0.89 |
| Pperoxisome | +1.03 | 0.584 | -1.79 | 0.002 | 6.1% | 93.9% | -0.95 | -1.07 | -1.03 | -0.95 |
| Allograft Rejection | +1.07 | 0.398 | -1.06 | 0.314 | 2.8% | 97.2% | -0.70 | -0.90 | +1.18 | -0.98 |

| Hallmark Pathway | Bulk NESperm | Bulk FDR | snRNA NESperm | snRNA FDR | Comp. % | Within-Cell % | Malignant NES | CAF NES | Vascular NES | CD8+ T NES |
|---|---|---|---|---|---|---|---|---|---|---|
| Spermatogenesis | +1.07 | 0.483 | -0.90 | 0.556 | 11.2% | 88.8% | -0.87 | -1.02 | -0.84 | -0.93 |
| KRAS Signaling Up | +1.28 | 0.006 | -1.42 | 0.017 | 3.0% | 97.0% | -0.79 | -0.99 | -1.08 | +1.04 |
| KRAS Signaling Dn | +0.98 | 0.764 | +1.01 | 0.084 | 75.1% | 24.9% | -0.91 | +0.25 | -0.96 | +1.14 |
| Pancreas Beta Cells | +1.13 | 0.398 | +2.07 | 0.002 | 88.8% | 11.2% | -0.72 | +1.19 | +0.92 | +1.26 |

*Note: NESperm values were computed via 1,000-permutation pre-ranked GSEA. Global cross-modal rank concordance: Spearman $\rho$ = 0.512 ($p = 1.46 \times 10^{-4}$), Pearson $r = 0.195$ ($p = 0.175$).*

**Supplementary Table S6. Summary of NRF2 signature correlations, Radiation Sensitivity Index (RSI) associations, TCGA-PAAD overall survival modeling, and single-nucleus sub-lineage NFE2L2 pseudobulk tests.**

| Analysis / Cohort | Comparison / Signature | Sample Size | Primary Metric | Exact Statistic / CI / p-value |
|---|---|---|---|---|
| Signature Correlation (TCGA-PAAD) | NRF2 Core Machinery vs. Iron/Ferroptosis Defense | n = 178 | Pearson r / Spearman ρ | $r = 0.569$ ($p = 1.11 \times 10^{-16}$); $\rho = 0.276$ |
| Signature Correlation (GSE225767) | NRF2 Core Machinery vs. Iron/Ferroptosis Defense | N = 55 | Pearson r / Spearman ρ | $r = 0.668$ ($p = 2.54 \times 10^{-8}$); $\rho = 0.716$ |
| RSI Correlation (TCGA-PAAD) | 8-gene NRF2 Target vs. 10-gene RSI | n = 178 | Pearson r / Spearman ρ | $r = 0.869$ ($p = 1.15 \times 10^{-55}$); $\rho = 0.786$ ($p = 1.33 \times 10^{-38}$) |
| RSI Correlation (GSE225767) | 8-gene NRF2 Target vs. 10-gene RSI | N = 55 | Pearson r / Spearman ρ | $r = 0.743$ ($p = 8.28 \times 10^{-11}$); $\rho = 0.669$ ($p = 2.36 \times 10^{-8}$) |
| Pathway Enrichment (TCGA-GTEx) | Reactome KEAP1-NFE2L2 (R-HSA-9755511) | n = 345 | ZGSEA / FDR | ZGSEA = +8.40, FDR < $1.71 \times 10^{-5}$ (59/95 leading-edge) |
| Pathway Enrichment (GSE225767) | GO Cellular Response to Oxidative Stress (GO:0034599) | N = 55 | ZGSEA / nominal p / FDR | ZGSEA = +2.07, nominal p = 0.0195, FDR = 0.131 (54/116) |
| Survival - Continuous Cox (TCGA) | 6-gene NRF2 Antioxidant Signature | N = 177 | Cox HR / Wald p / PH test | HR = 1.35 (95% CI: 1.01-1.81), p = 0.0422, PH p = 0.690 |
| Survival - Exploratory Cutpoint (TCGA) | 6-gene NRF2 (135 High vs. 42 Low) | N = 177 | Median OS / Log-rank / maxstat | 19.6 vs. 21.7 mo; unadj. p = 0.0462, maxstat padj = 0.393, HR = 1.60 |
| Survival - Continuous Cox (TCGA) | 3-gene Iron/Ferroptosis (FTH1, FTL, GPX4) | N = 177 | Cox HR / Wald p / PH test | HR = 1.07 (95% CI: 0.82-1.39), p = 0.630, PH p = 0.872 |

| Analysis / Cohort | Comparison / Signature | Sample Size | Primary Metric | Exact Statistic / CI / p-value |
|---|---|---|---|---|
| Survival - Exploratory Cutpoint (TCGA) | 3-gene Iron/Ferroptosis (67 High vs. 110 Low) | N = 177 | Median OS / Log-rank / maxstat | 17.0 vs. 21.7 mo; unadj. p = 0.0908, maxstat padj = 0.590, HR = 1.45 |
| snRNA Pseudobulk (GSE202051 All) | NFE2L2: Neoadjuvant (n = 25) vs. Naïve (n = 18) | N = 43 | BH FDR q / nominal p | ↑ Dendritic, Delta, Vascular (q = 0.027); ↑ Alpha (q = 0.047); ↓ Malignant (p = 0.02) |
| snRNA Pseudobulk (GSE202051 CRT) | NFE2L2: Standard CRT (n = 14) vs. Naïve (n = 18) | N = 32 | Expressing fraction / shifts | Malignant nuclei ~45% → ~35%; ↓ Malignant/Ductal/B/VSMC; ↑ Pericyte/Vascular |
