## Supplementary figures for "PDAC BioPortal: An Integrated Web Resource for Multi-Scale Analysis of Pancreatic Cancer Transcriptomics"

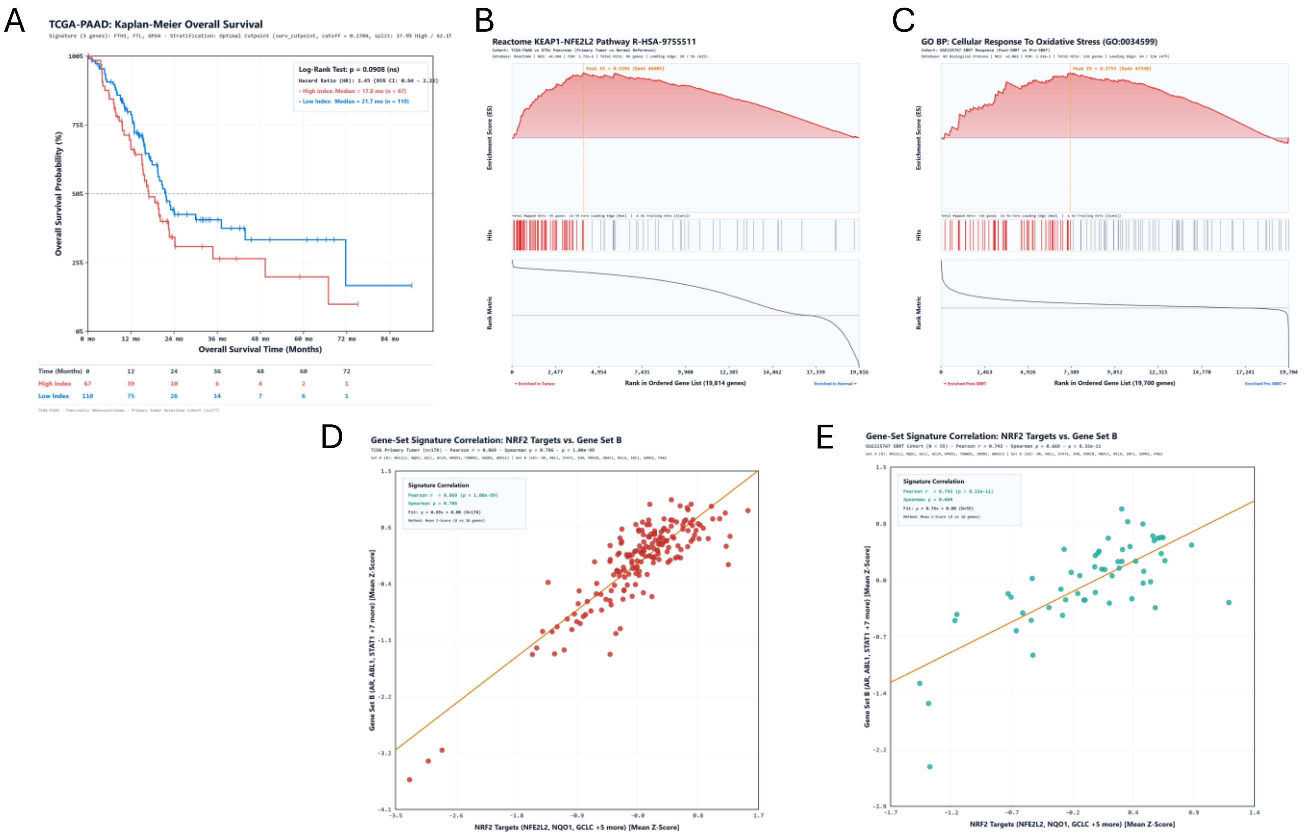


**Supplementary Figure S1. Pathway-level NRF2 enrichment, survival association of ferroptosis-defense genes, and correlation with the Radiation Sensitivity Index (RSI).** (**A**) Kaplan-Meier overall survival analysis in TCGA-PAAD ($N$ = 177) stratified by exploratory optimal cutpoint of the three-gene Iron & Ferroptosis Defense signature (*FTH1*, *FTL*, *GPX4*; High $n$ = 67, median OS 17.0 months vs. Low $n$ = 110, median OS 21.7 months; unadjusted log-rank $p$ = 0.0908, Miller-Siegmund maxstat $p_{adj}$ = 0.590, HR = 1.45, 95% CI: 0.94-2.22; continuous univariate Cox HR = 1.07 per unit $Z$-score, 95% CI: 0.82-1.39, Wald $p$ = 0.631). (**B**) GSEA enrichment plot for Reactome KEAP1-NFE2L2 signaling (R-HSA-9755511) in TCGA-PAAD primary tumors vs. GTEx normal pancreas ($Z_{GSEA}$ = +8.40 [portal UI NES], FDR < 1.71 × $10^{-5}$; 59/95 leading-edge hits). (**C**) GSEA enrichment plot for GO Biological Process cellular response to oxidative stress (GO:0034599) in GSE225767 post-SBRT vs. pre-SBRT samples ($Z_{GSEA}$ = +2.07 [portal UI NES], nominal $p$ = 0.0195, FDR = 0.131; 54/116 leading-edge hits). (**D-E**) Sample-level correlation between an eight-gene NRF2 target signature (*NFE2L2*, *NQO1*, *GCLC*, *GCLM*, *HMOX1*, *TXNRD1*, *SRXN1*, *AKR1C1*) and the 10-gene Radiation Sensitivity Index (RSI; *AR*, *ABL1*, *STAT1*, *JUN*, *PRKCB*, *HDAC1*, *RELA*, *IRF1*, *SUMO1*, *PAK2*) in (**D**) TCGA primary PDAC tumors ($n$ = 178; Pearson $r$ = 0.869, $p$ = 1.15 × $10^{-55}$; Spearman $\rho$ = 0.786, $p$ = 1.33 × $10^{-38}$; note that the interactive portal panel displays $p$ = 1.00 × $10^{-99}$ due to UI floor clamping below $10^{-50}$) and (**E**) the GSE225767 SBRT cohort ($N$ = 55; Pearson $r$ = 0.743, $p$ = 8.28 × $10^{-11}$; Spearman $\rho$ = 0.669, $p$ = 2.36 × $10^{-8}$).

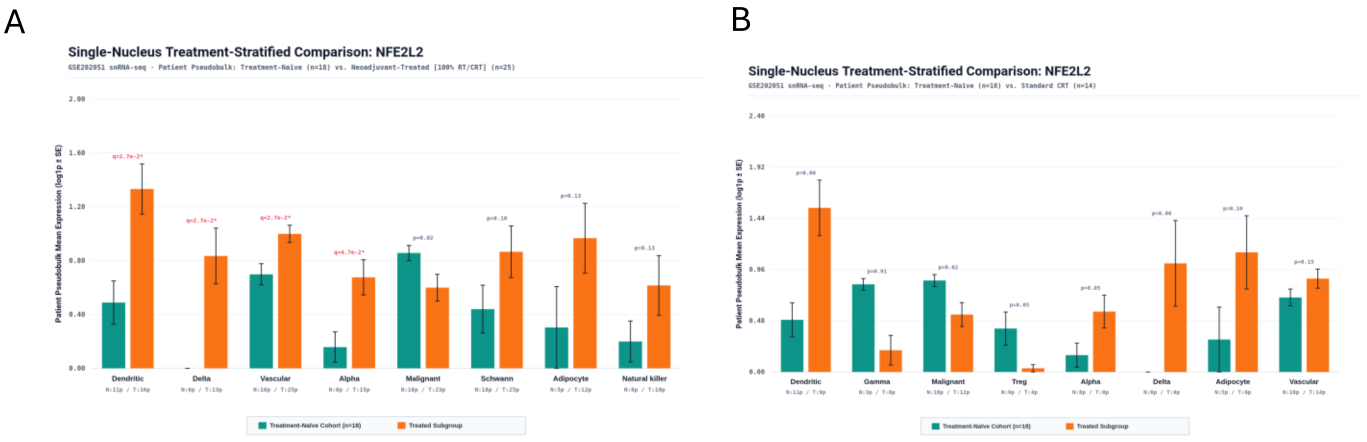


**Supplementary Figure S2. Patient-level pseudobulk comparison of *NFE2L2* expression across PDAC cellular sub-lineages.** Mean patient-level pseudobulk *NFE2L2* expression ($\log_{1p}$ ± SE) comparing treatment-naïve patients ($n$ = 18, teal) against (**A**) the full neoadjuvant-treated cohort ($n$ = 25, orange; FDR $q$ = 0.027 for dendritic, delta, and vascular endothelial cells; $q$ = 0.047 for alpha cells; nominal $p$ = 0.02 for malignant epithelial cells) and (**B**) the standard chemoradiotherapy (CRT) subgroup ($n$ = 14, orange). Patient sample counts per compartment ($N$/$T$) are indicated below each sub-lineage.

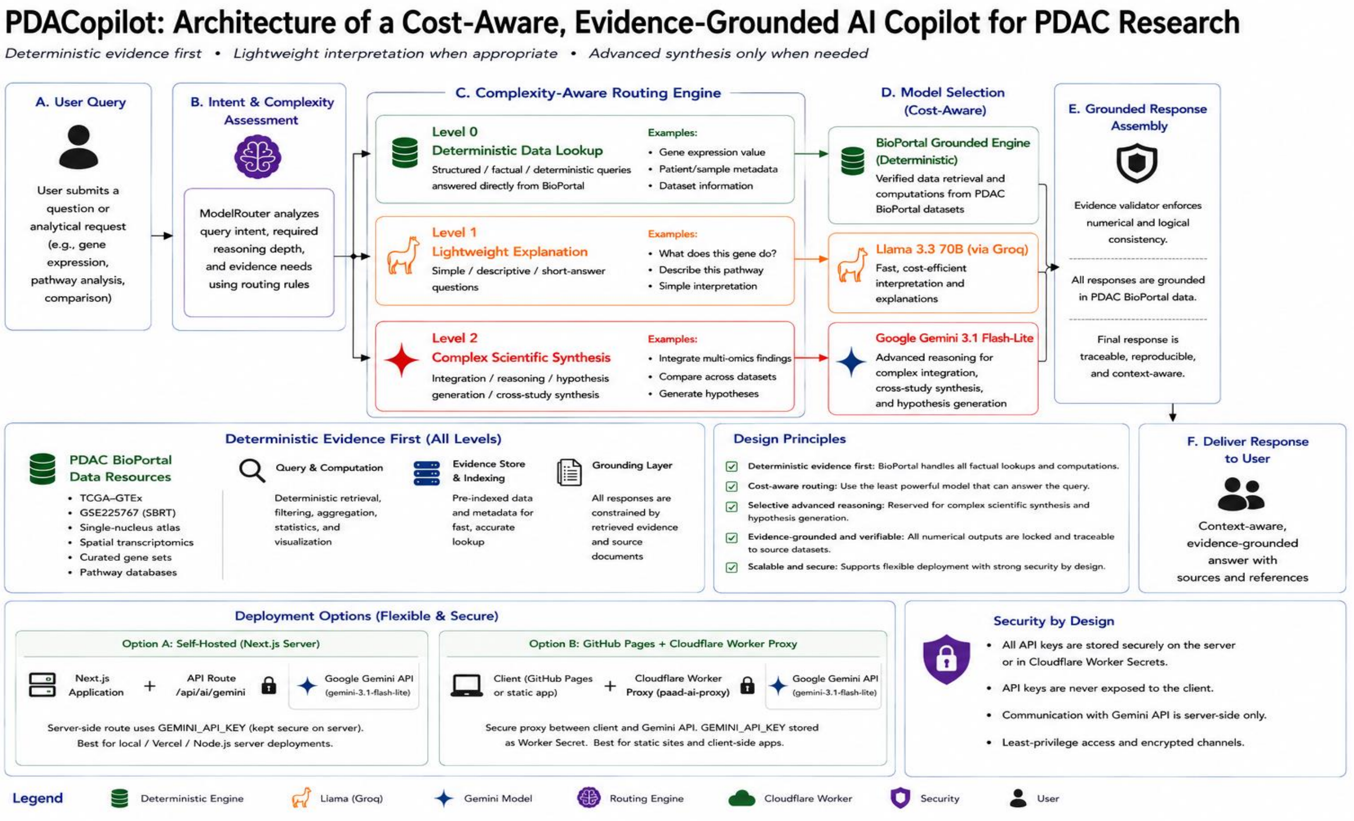


**Supplementary Figure S3. Cost-aware, evidence-grounded hierarchical routing architecture of PDACopilot.** (**A-F**) Execution pipeline from natural-language user query (**A**) and ModelRouter intent/complexity classification (**B**) through three hierarchical resolution tiers (**C-D**): Level 0 (Deterministic BioPortal Grounded Engine), Level 1 (Llama 3.3 70B lightweight interpretation), and Level 2 (Google Gemini Flash complex cross-modal synthesis), followed by EvidenceValidator consistency gating (**E**) and delivery of traceable, dataset-cited responses (**F**). Lower panels summarize deterministic evidence grounding, design principles, deployment topologies (Next.js server API route and Cloudflare Worker proxy), and API key security.
